# Clearing of ribosome collisions by the ribosome quality control trigger complex RQT

**DOI:** 10.1101/2022.04.19.488791

**Authors:** Katharina Best, Ken Ikeuchi, Lukas Kater, Daniel Best, Joanna Musial, Yoshitaka Matsuo, Otto Berninghausen, Thomas Becker, Toshifumi Inada, Roland Beckmann

## Abstract

After translational stalls, colliding eukaryotic ribosomes are cleared through dissociation into subunits by the ribosome quality control trigger complex, RQT, by an unknown mechanism. Here we show that RQT requires accessible mRNA and the presence of a neighboring ribosome. Cryo-EM of several RQT-ribosome complexes revealed the structural basis of splitting: RQT engages the 40S subunit of the lead ribosome and can switch between two conformations. We propose a mechanistic model in which the Slh1 helicase subunit of RQT applies a pulling force on the mRNA, causing destabilizing conformational changes of the 40S subunit. The collided ribosome functions as a ram or giant wedge, ultimately resulting in subunit dissociation. Our findings provide a first conceptual framework for a helicase driven ribosomal splitting mechanism.

**One-Sentence Summary:** RQT clears collided ribosomes by pulling mRNA to trigger destabilizing conformational transitions for subunit dissociation.

## Introduction

Ribosomal stalling can occur during the translation of messenger RNAs. While transient ribosome stalls may fulfill biological functions, e.g. for co-translational protein folding or targeting, persistent stalls usually indicate a stress situation that requires a response. Such persistent stalling events result in the collision of trailing ribosomes with the stalled ribosome, thereby creating new composite interfaces which are then recognized by cells as a proxy for translational stress. Accordingly, these collisions were found to be interaction hubs for numerous factors mediating different translation-based control pathways, such as the integrated stress response (ISR), the ribotoxic stress response (RSR) or the ribosome-associated quality control (RQC) pathway (*1-3*).

The RQC pathway is triggered after such stalling on rare codons or poly-A stretches, and results in abortion of translation of the aberrant mRNA, recycling of the stalled ribosome and degradation of the arrested potentially toxic protein product by the ubiquitin-proteasome system (*4-6*). Upon ribosome collision RQC is initiated by (poly-)ubiquitination of several ribosomal proteins near the mRNA entry channel by the E3 ligase Hel2 (ZNF598 in mammals) (*7-13*). This is followed by disassembly of the ubiquitinated collided ribosomes by the ribosome quality control triggering complex (RQT) in yeast (ASC-1 or hRQT complex; ASCC in humans) (*7, 8, 14, 15*) which was shown to first disassemble the lead (stalled) ribosome of the ribosome queue (*8*). This event generates a 60S subunit still attached to a peptidyl-tRNA, that is the target of the downstream ribosome-associated quality control (RQC) pathway. Here, in a template-less peptide synthesis event, the RQC-complex adds C-terminal alanine/threonine tails (CAT-tails) via its Rqc2 subunit (*16*), whereas the E3 ligase Ltn1 poly-ubiquitinates the nascent peptide for degradation (*17-19*).

The RQT complex of yeast is composed of the ATP-dependent Ski2-like RNA helicase Slh1 (Rqt2), the ubiquitin-binding CUE domain containing protein Cue3 (Rqt3) and the zinc-finger containing protein Rqt4 (*8*). The main ribosome splitting activity has been attributed to the helicase Slh1, however, the underlying mechanism of ribosome disassembly by RQT has remained largely enigmatic. The requirement of Hel2-dependent ubiquitination of uS10 and ATP hydrolysis has been shown (*8*), yet, it is unclear how RQT interacts with collided ribosomes and how it employs its RNA helicase activity for selective splitting of the stalled lead ribosome in a ribosome queue.

## Results

### Processive splitting by RQT is dependent on accessible mRNA and a trailing 80S

We set out to characterize the helicase dependent splitting activity of RQT biochemically. The closest homolog of its Slh1 (Ski2-like helicase 1) helicase subunit is Brr2 (bad response to refrigeration 2), a tandem-helicase cassette containing protein that interacts with single stranded RNA to unwind base-paired spliceosomal U4:U6 snRNA for spliceosome activation (*20*). The eponymous Ski2 helicase was shown to bind to mRNA emerging from ribosomes and catalyze the extraction of the mRNA for subsequent mRNA decay by the cytoplasmic exosome (*21-23*). Assuming that the Slh1 helicase (Fig. 1A), also engages mRNA and considering that RQT is known to split the first/stalled ribosome, Slh1 would require the presence of an accessible 3’ mRNA overhang emerging from the stalled lead ribosome.

**Fig. 1:**
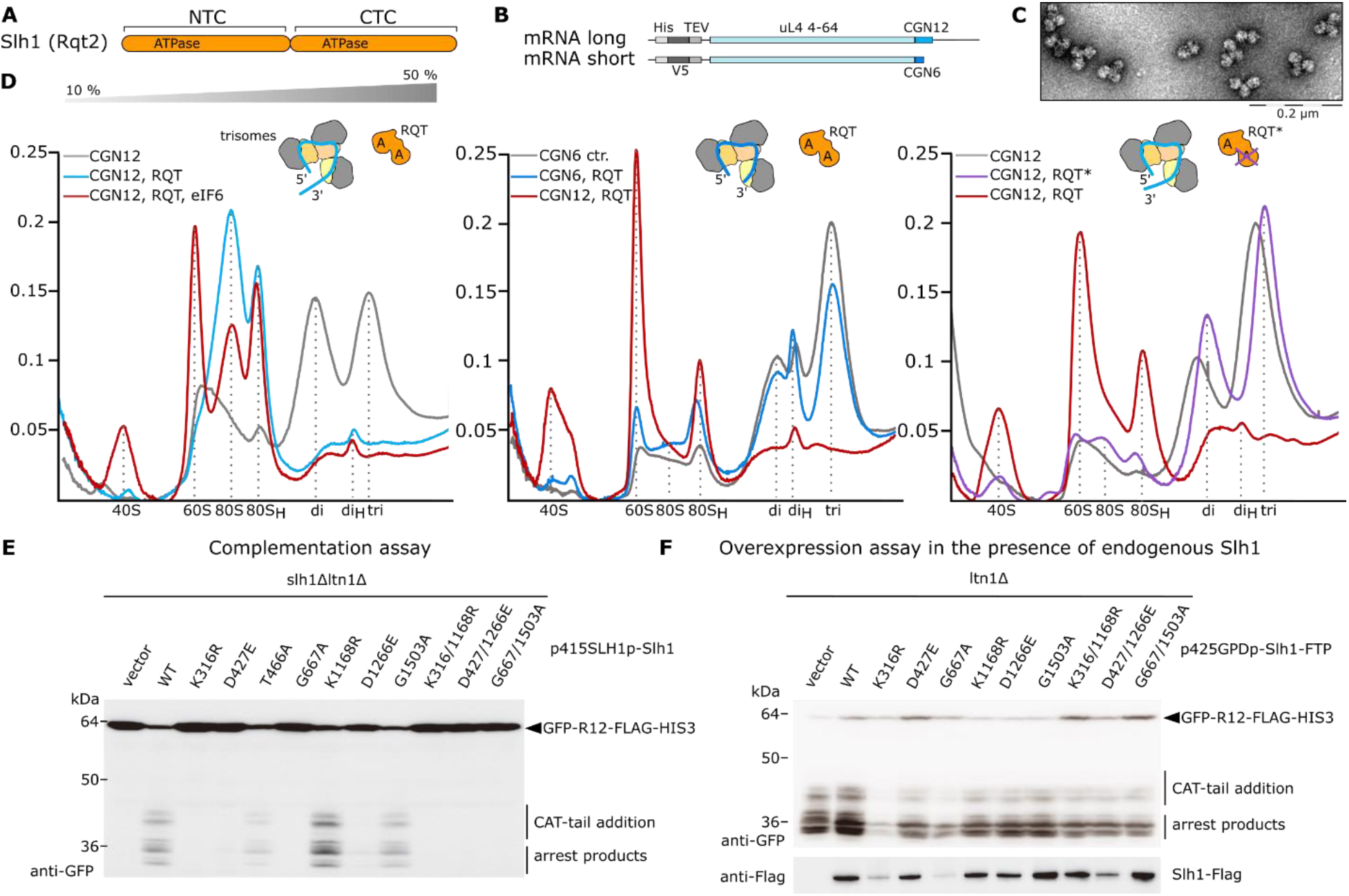
Biochemical analysis of RQT-dependent splitting of collided ribosomes. (**A**) Scheme outlining the tandem ATPase cassettes of Rqt2 (Slh1). NTC, CTC = N-terminal and C-terminal cassette; (**B**) Scheme outlining the mRNA constructs used for generating (hyper-)ubiquitinated trisomes. CGN6 and CGN12 = stalling mRNA sequence contain 6 or 12 consecutive CGN (N=A, G or C) codons. (**C**) Uranyl-actetate stained negative stain-TEM image of polyubiquitinated trisomes. (**D**) Polyribosome profiles from sucrose density gradients obtained after *in vitro* splitting reactions. Left: Splitting reactions using trisomes with a long 3’-mRNA stretch emerging from the mRNA entry site of the lead ribosome (stalled on CGN_12_ mRNA) and of RQT with intact helicase activity (see cartoon representation; A = ATPase). Grey, trisomes only; blue, splitting reaction without and red, with eIF6. 80_H_ = 80S-40S halfmers; di_H_ = 40S-disome halfmers. Middle: Comparison of splitting reactions using trisomes with long versus short stretches of 3’-mRNA (stalled on CGN_6_ mRNA); right: Comparison of splitting reactions using wild type versus helicase deficient K316R-RQT (RQT*). (E, F) *CGN12* reporter gene assay to monitor RQC activity in *SLH1* mutants: (**E**) Complementation analysis: *SLH1* wild-type and the mutants were expressed by endogenous *SLH1* promoter in the *slh1Δltn1Δ* double deletion mutant. (**F**) Overexpression analysis: *SLH1* wild-type and the mutants were highly expressed by *GPD* (*TDH3*) promoter in the *ltn1Δ* deletion mutant. The products derived from *GFP-CGN(R)12-FLAG-HIS3* reporter gene were monitored by immunoblotting using anti-GFP antibody.

To test this, we optimized our previously described yeast *in vitro* system (*8*) to reconstitute and monitor RQT-mediated splitting of ubiquitinated collided ribosomes (Suppl. Fig. 1A) with different mRNA constructs. In brief, a cell-free *in vitro* translation extract was generated from an mRNA stabilizing yeast strain (Δ*cue2*, Δ*slh1*, Δ*xrn1*) in order to stably stall ribosomes on mRNA reporters containing stretches of hard-to-decode CGN (N= A, C or G) codons (Fig. 1B, Suppl. Fig. 1B). These mRNA reporters were designed such that after stalling at the second or third CGN codon in the P-site (*7*) 3’-mRNA regions of different length emerge from the mRNA entry of the stalled lead ribosome: in the (CGN)_12_ control construct containing twelve CGN codons and an additional 3’-mRNA region, an overhang of about 150 nt emerges from the ribosome and would be accessible to RQT, whereas in the constructs with shorter stall repeats, (CGN)_6_ and (CGN)_4_ and no additional 3’ region, only a 9-12 nt or a 3-6 nt long 3’ stretch follows the A-site, and is too short to emerge from the ribosome and could thus not serve as a substrate for RQT.

After translation of these mRNA constructs and subsequent affinity purification of ribosome-nascent chain complexes (RNCs), di- and trisomes were harvested from a sucrose density gradient (Suppl. Fig. 1C), quality controlled by Negative Stain-TEM (Fig. 1C), and *in vitro* poly-ubiquitinated using purified Hel2 (E3), Ubc4 (E2), Uba1 (E1), ubiquitin and ATP (Suppl. Fig. 1D, 1E). Then, splitting reactions with the ubiquitinated trisome fraction were performed in the presence of ATP, a molar excess of purified wild type RQT (Suppl. Fig. 1E) or RQT with an ATPase-deficient Slh1-K316R mutant (*8*) (K316R-RQT), as well as purified anti-association factor eIF6. Subsequently, the reactions were analyzed by sucrose density gradient centrifugation.

Reactions with ubiquitinated (CGN)_12_ stalled trisomes containing the long accessible mRNA 3’-region resulted in a near complete collapse of the trisome and also disome peak, accompanied by an increase of 40S subunit, 60S subunit and 80S monosome peaks (Fig. 1D, left panel). The control reactions of the same trisomes in the absence of RQT and ATP displayed only a stable trisome peak and an additional disome peak (Fig. 1D), probably as a result of unspecific background nuclease or other dissociation activity. This showed that the RQT complex, as expected, efficiently resolves the collisions by splitting of ribosomes into subunits. Yet, in contrast to the efficiently split (CGN)_12_ trisomes, the splitting activity was largely abolished when using trisomes with the very short 3’ region of the (CGN)_6_ or the (CGN)_4_ constructs with a minimal accessible 3’ mRNA overhang (Fig. 1D, middle panel and Suppl. Fig. 1F).

This indicated that the RQT activity depends on freely accessible mRNA and supports our hypothesis that the Slh1 helicase activity acts on the mRNA and not the rRNA. Splitting activity was also lost when using trisomes without prior *in vitro* ubiquitination (Suppl.Fig 1G) or when using an ATPase-deficient mutant RQT(K316R) (Fig. 1D, right panel), consistent with previous studies using *SDD1* mRNA as a reporter (*8*). In addition, Walker mutations in both the N- and the C-terminal helicase cassettes (NTCs and CTCs) showed severe effects on the splitting-dependent CAT tailing reaction using a CGN_12_ stalling reporter *in vivo*. This was observed in complementation assays in which a *SLH1* knockout strain was probed for CAT tailing when complemented with different Slh1 constructs (Fig. 1E). Overexpression of Slh1 constructs in the presence of the wildtype gene revealed that two of the NTC mutations display a dominant negative phenotype for CAT tailing (Fig. 1F). Together, this suggested that the activity of both ATPases is required to engage the emerging mRNA and trigger efficient splitting.

We noticed that the second ribosome of a queue was efficiently split in our reaction, most likely following the dissociation of the first ribosome in a processive manner. This was indicated by the almost complete disappearance of not only the trisome but also the disome peak. Importantly, however, the last trailing 80S was not split by RQT since we observed accumulation of 80S ribosomes. This notion is further supported by the presence of 40S-80S and 40S-80S-80S halfmer peaks, likely representing splitting products in which, after dissociation of the 60S of the lead ribosome, the 40S is still connected with trailing ribosomes via the 40S-40S collision interface or via its mRNA association. In order to corroborate that 80S ribosomes without a collided neighbor ribosome are poor substrates for the RQT complex, we performed the splitting assay with a preparation containing mainly isolated ubiquitinated 80S ribosomes and observed that these individual 80S monosomes were at most modestly dissociated by RQT (Suppl. Fig. 1H). This suggests that the RQT-dependent splitting mechanism requires a neighboring (trailing) collided ribosome which is in line with our observation that the *in vitro* trisome splitting reactions do not target the last 80S ribosome.

Taken together we conclude that (i) RQT splits with its Slh1 helicase component exerting force primarily on the mRNA, specifically by engaging the 3’ region of the mRNA emerging from the lead ribosome, (ii) that RQT splits in a processive manner and (iii) that RQT requires at least one neighboring ribosome for efficient splitting.

### Cryo-EM analysis of RQT-bound ribosomes in pre- and post-splitting reactions

To obtain deeper mechanistic insights into the splitting process, we performed cryo-electron microscopy (cryo-EM) analysis of our RQT-splitting reactions (Suppl. Fig. 2-6, Table S1). To limit the apparent complexity of the trisome sample (*8*) we used the *in vitro* ubiquitinated disomes instead of trisomes. We flash-froze the samples quickly after addition of RQT in an attempt to capture reaction intermediates. We expected that reactions containing ubiquitinated disomes and the mutant K316R-RQT or disomes without accessible mRNA overhang ((CGN)_6_-disomes) would enrich pre-splitting intermediates, whereas reactions using the disomes with a long mRNA overhang ((CGN)_12_-disomes) and wild-type RQT would enrich mainly post-splitting states.

**Fig. 2:**
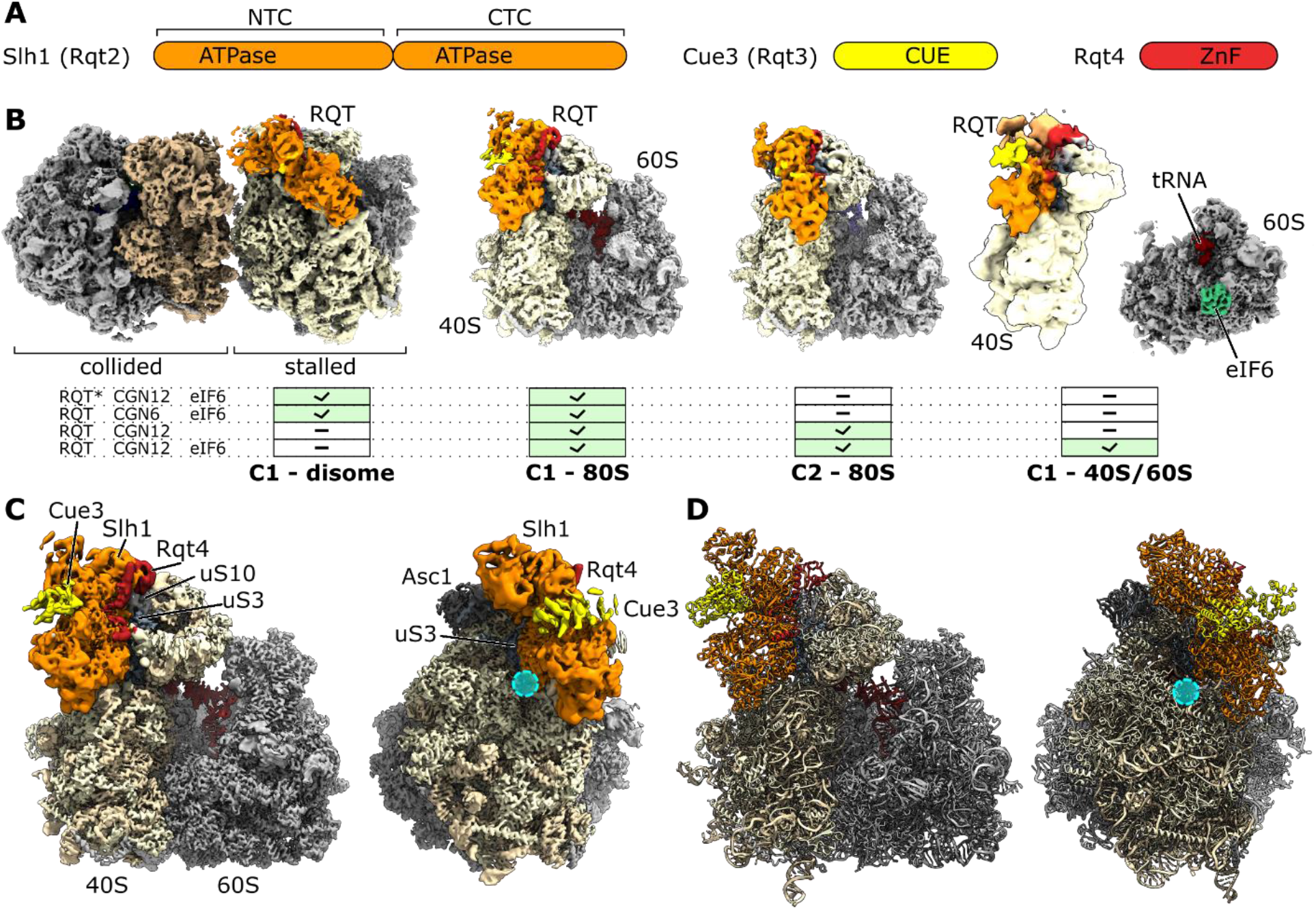
Cryo-EM analysis of RQT-ribosomes complexes. (**A**) Scheme outlining components of the trimeric RQT complex. NTC, CTC = N-terminal and C-terminal cassette; CUE = Coupling of Ubiquitin to ER degradation domain; ZnF = zinc-finger domain. (**B**) Composite cryo-EM maps of RQT-containing ribosomal particles found in cryo-EM analyses of pre-splitting reactions (short mRNA or ATPase deficient Slh1, RQT*) and post-splitting (long mRNA and wt Slh1) reactions. Results are summarized in a table. C1 and C2 = two different conformations of ribosome-bound RQT. (**C, D**) Cryo-EM map (C) and molecular model (D) of the RQT-80S in C1 conformation. All maps are shown as composite maps after local refinement (see Suppl. Figs 2-5 for details). The mRNA entry site is highlighted with a cyan circle.

Extensive 3D classification (Suppl. Fig. 2-3) of pre-splitting samples with short mRNA and ATPase deficient Slh1 showed a large population of intact disomes in their canonical conformation (*11, 13*) prior to splitting: the lead ribosome adopted a non-rotated post-translocational (POST) state 80S with a P/P tRNA, whereas the trailing ribosome adopted a rotated PRE-state with hybrid A/P-P/E tRNAs. A subpopulation of these disomes was associated with additional density which we could unambiguously assign to the RQT complex with all three subunits (Fig. 2A, 2B, left). The complex was located on the 40S subunit in close proximity to the mRNA entry tunnel and the ribosomal protein uS10 which has been shown to carry the poly-ubiquitination modification that signals RQT recruitment (*7, 8*). While we did not observe density for this modification in any of our structures (probably due to its dynamic nature), the close vicinity between uS10 and RQT is in line with direct interaction licensing RQT for ribosome binding. We found RQT exclusively on the lead but not on the trailing ribosome, in agreement with previous biochemical findings that RQT preferentially splits the lead stalled ribosome of a ribosome queue (*8*). This preference can be explained now since in the presence of the lead ribosome it would be impossible for the RQT complex to engage a trailing ribosome without resulting in severe steric clashes; its binding site on the 40S subunit is almost completely masked by the lead ribosome (Suppl Fig. 7A). After completed dissociation of the first ribosome, however, RQT could easily engage the next ribosome in a collided queue, explaining the observed sequential activity on the trisome. We noticed that density for the RACK1/Asc1 protein of the lead ribosome found at the collision interface (*8, 11, 13*) was underrepresented, probably indicating that destabilizing forces are already in place simply upon ribosome binding of RQT to the collision (Suppl. Fig. 7B, 7C).

In addition to disomes, these samples contained 80S monosomes, most likely as a result of background dissociation activity; these 80S monosomes adopted the POST-state with RQT bound in the same position on the 40S subunit as in the disome (Fig. 2B). Yet, no small or large subunits were found when using the ATPase-dead mutants despite the presence of eIF6 in these samples, consistent with the lack of splitting activity observed in the corresponding *in vitro* assays.

In contrast, when analyzing the samples treated with wt ATPase activity of RQT, we mainly observed post-splitting products with almost no stable disomes being detected anymore (Suppl. Figs 4 and 5). The majority of particles represented various populations of 80S monosomes and a substantial amount of 40S subunits as well as 60S subunits carrying a peptidyl-tRNA and eIF6 (Fig. 2B), together confirming that splitting reactions took place. Also here, density for the RQT complex was present on 80S monosomes, but contrary to the pre-splitting situation, we found these 80S ribosomes and their associated RQT in two different conformations: the classical POST state of the lead ribosome (termed C1) (Fig. 2B, 2C) and a state termed C2 in which the 40S head was swiveled by 20 degrees relative to the 40S body and RQT was repositioned on the 40S (Fig. 2B). This C2 state was present in both eIF6-free but also eIF6-containing datasets indicating that it is not a result of re-association. Moreover, in the presence of eIF6, we also observe a 40S population with RQT bound in the C1 state conformation which may represent the expected splitting product before dissociation of the RQT complex (Fig. 2B). High-resolution refinement of the RQT-bound ribosome classes followed by local refinement of the isolated RQT densities (Suppl. Fig. 6) allowed us to build molecular models for RQT-ribosome complexes based initially on AlphaFold2 (*24*) predictions for the RQT subunits Slh1, Cue3 and Rqt4 (Suppl. Fig. 8 and Suppl. Fig. 9). Overall, these models reveal the overall architecture of the RQT and how it interacts with the head and body of the 40S subunit in close proximity to the mRNA entry tunnel (Fig. 2D and Fig. 3).

**Fig. 3:**
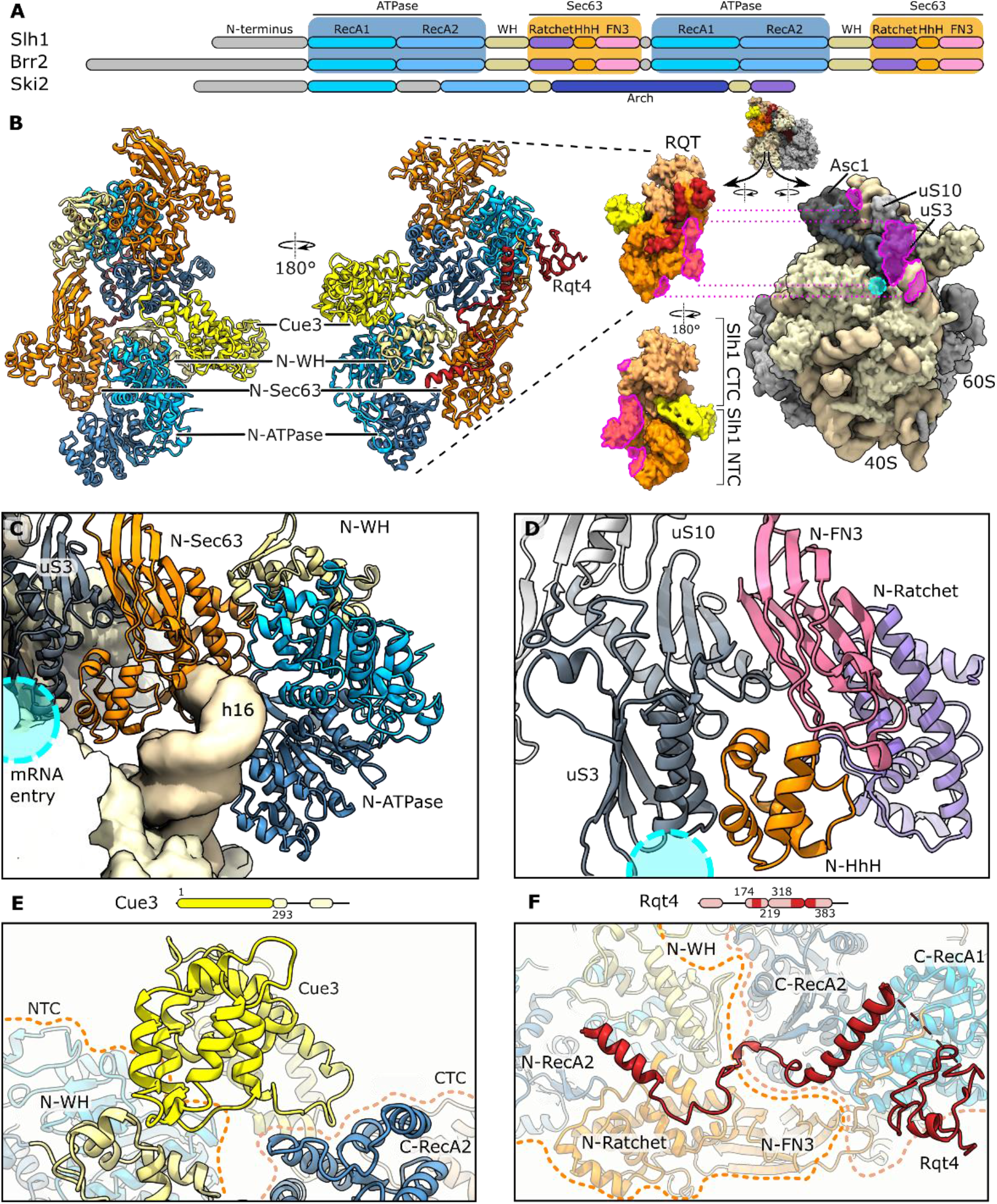
Molecular model of ribosome-bound RQT. (**A**) Domain arrangement of Slh1 and closest homologs Brr2 and Ski2. (**B**) Molecular model of RQT bound to the 80S ribosome in C1 state. On the right side RQT and the 80S ribosome were converted to low-pass filtered density for clarity. RQT-ribosome interaction regions are highlighted in violet. The mRNA entry site is marked as a blue circle. A thumbnail indicates the overall orientation. (**C**) View focusing on the interaction of Slh1-NTC with the 40S. (D) Zoom view showing the interaction of the N-terminal Sec63 homology region of Slh1 with 40S uS3. (**E**) View focusing on the Cue3 N-terminal (helical) domain interacting with Slh1. (**F**) View focusing on Rqt4 interacting with Slh1. In (**E**) and (**F**) schematic representations of Cue3 and Rqt4 are shown. Modelled residues are shown in strong, unmodeled regions in weak color; dashed orange and pale, yellow lines outline N- and C-terminal cassettes (NTC and CTC) of Slh1.

**Fig. 4:**
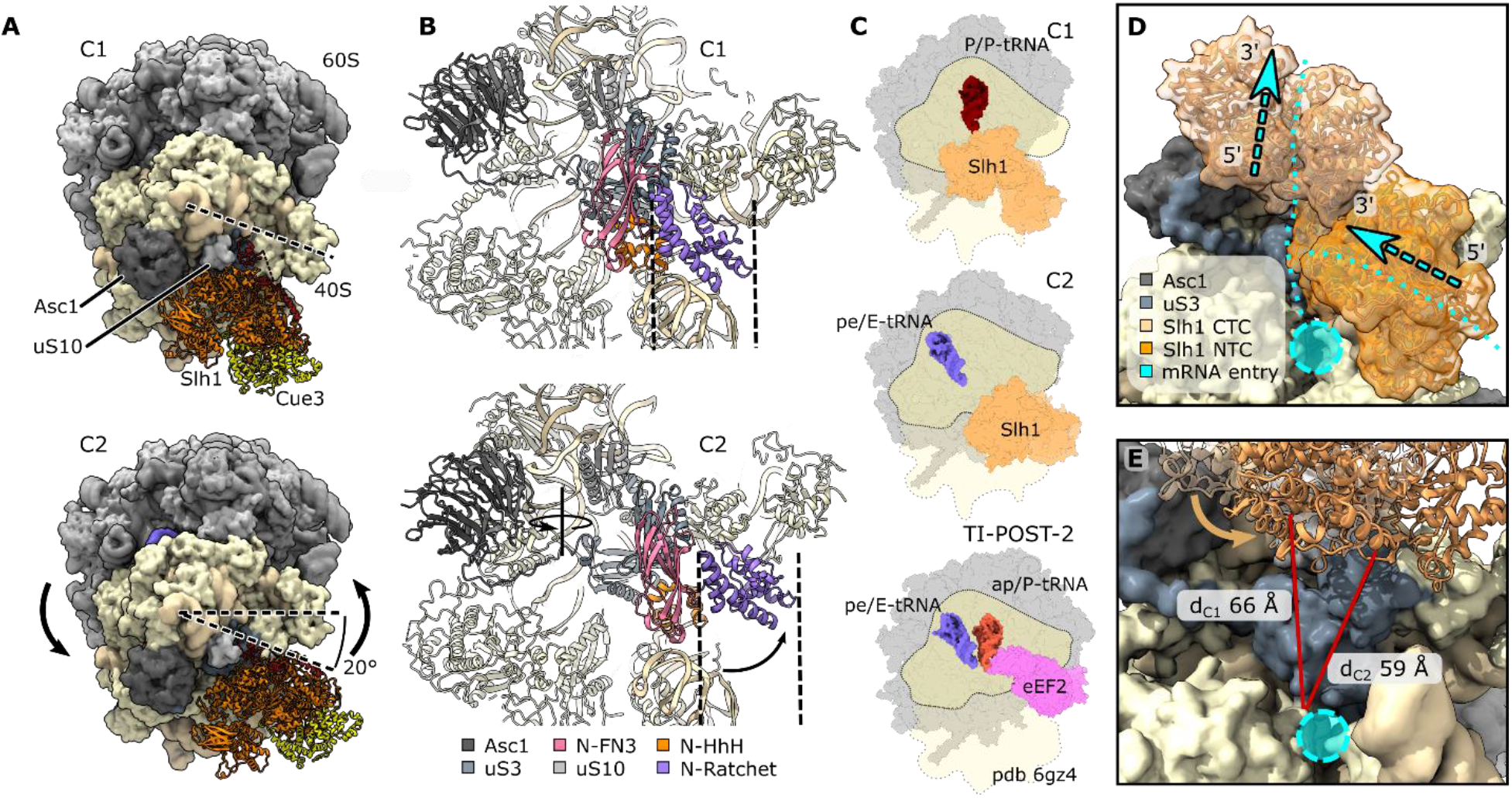
Conformational dynamics of the RQT-ribosome complexes. (**A, B**) Comparison of C1 and C2 states of the RQT-80S complex. (**A**) Top views; in C1, the dashed line marks the position of the 40S head; in C2, the rotation direction and resulting net rotation angle for the head-swivel movement is indicated (20°). For clarity the molecular model of the ribosome was converted to low-pass filtered density. (**B**) View focusing on the main 40S contacts of the RQT NTC and its repositioning. The solid line in (**B**) displays the rotation axis of the 40S head swivel; RQT undergoes an additional movement and dashed lines assist in assessing the shift of RQT domains relative to the invariant 40S body. (**C**) Top views of cartoon representations comparing Slh1-bound C1 and C2 80S complexes with the TI-POST-2 translocation intermediate as observed in the eEF2-GMP-PNP-bound mammalian 80S (*29*). (**D**) Zoom view focusing on the relative position of Slh1’ helicase cassettes on the mRNA entry site. Arrows indicate the directions of movement of the 3’-5-helicase and the mRNA. (**E**) View focusing on the location of Slh1 CTC in C1 (transparent ribbons) and C2 (solid ribbons) states above the mRNA entry channel. Red lines indicate the distane between the helicase entry site and an invariant residue (uS5) of the 40S body nearby the mRNA entry site (cyan circle). Note that Slh1 CTC is 7 Å closer to the mRNA entry site.

**Fig. 5:**
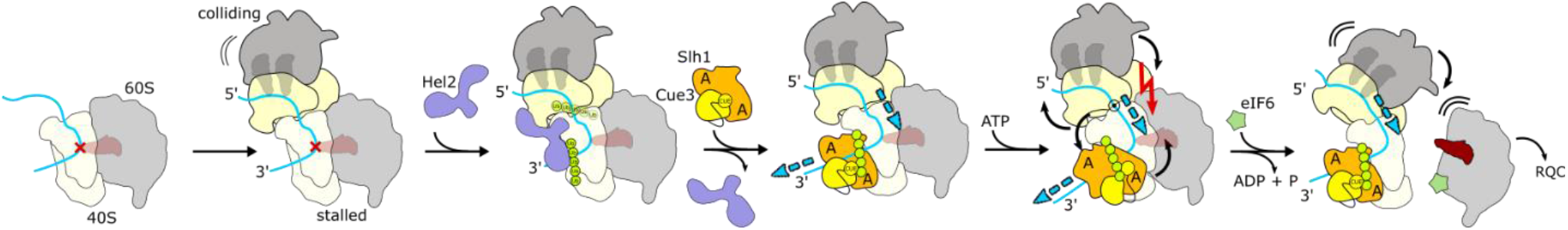
Model for RQT-mediated ribosome splitting after collision. After stalling and subsequent collision of trailing ribosomes, Hel2 recognizes the collision interface and (poly-)ubiquitinates uS10. RQT binds polyubiquitinated stalled ribosomes and engages with accessible 3’-mRNA emerging from the stalled lead ribosome. A pulling force on the mRNA (blue arrows) may cause head-swiveling, destabilization of the lead ribosome and ultimately drive the trailing ribosome like a wedge between the subunits of the lead ribosome. A = ATP-binding site See text for details.

### Structure of the ribosome-bound RQT complex

The largest subunit of RQT, the Slh1 helicase contains, similar to its closest relative Brr2 (*25*), a helical N-terminal domain and a duplicated pair of RecA-like ATPase domains (RecA1 and RecA2), both of which are followed by a winged helix domain (WH) and a Sec63-like domain. The Sec63-like domain is subdivided into Ratchet-, helix-hairpin-helix (HhH) and fibronectin type III (FN3) domains. (Fig. 3A, Suppl. Fig. 8A). When bound to the ribosome, Slh1 adopted a bi-lobed arch-like architecture formed by the two RecA-WH-Sec63 cassettes (Fig. 3B, Suppl. Fig. 9B). The N-terminal cassette (NTC) is clamped between the body and the head at the mRNA entry site of the 40S subunit, stretching from helix 16 (h16) of 18S rRNA along the array of ribosomal proteins (r-proteins) eS10, uS3 and uS10 (Fig. 3B and 3C). The C-terminal cassette (CTC) is packed on the NTC and connects the RQT complex with RACK1 (Asc1) at the back side of the 40S head (Fig. 3B). Densities for both Cue3 and Rqt4 were positioned between the two cassettes with Cue3 binding to the upper convex surface and Rqt4 packing against the concave side of the Slh1 arch close to uS10 (Fig. 3B, 3E, 3F and Suppl. Fig.9B).

Our model for Slh1 accounts for the majority of the 1967 amino acid long protein, lacking only the N-terminal domain (1-217) (Suppl. Fig. 8A). The main anchor of Slh1 in both C1 and C2 conformations is constituted by part of the NTC Sec63-domain. Specifically, the HhH domain and the β-sandwich structure of the FN3 domain bind to uS3 at the inner side of the mRNA entry tunnel (Fig. 3C, 3D). The FN3 domain packs with its terminal β-sheet (which is absent in the CTC FN3 domain) against the β-sheet of the N-terminal uS3 KH domain; the HhH domain is anchored on the two antiparallel α-helices of the uS3 middle domain (Fig. 3D). In C1, the HhH forms additional contacts to eS30, another protein lining the mRNA entry path located in the 40S body near rRNA h16 (Suppl. Fig. 9E). This rRNA helix is a common interaction hub for various ribosome-interacting proteins (*26, 27*) including Ski2 (*22, 23*), and is accommodated in a deep pocket formed by the RecA2, Ratchet and HhH domains of the NTC (Fig. 3C). Finally, we observe extra density for the usually flexible uS5 N-terminus located on top of the Slh1 HhH/FN3 domain (Suppl. Fig. 9C).

Compared to the more compact structures of the Brr2 protein (Suppl. Fig. 10F), contacts between the two individual cassettes in Slh1 are limited to a few interactions between the WH domain and a part of the FN3 domain of the NTC and RecA2 of the CTC, forming a patch of mainly hydrophobic interactions (Suppl. Fig. 9C). Thus, Slh1 adopts a rather open, more elongated conformation when compared to Brr2 (Suppl. Fig. 10F). This allows the only direct interaction of the CTC with the 40S, established between its WH domain and RACK1 (Fig. 3B). Moreover, this distinct inter-cassette conformation is stabilized by Cue3 and Rqt4 (Fig. 3B, 3E and 3F, Suppl. Fig. 10H): Rod-like extra densities, wedged between the NTC’s WH domain and the CTC RecA2 at the concave side of Slh1, were assigned to the N-terminus (1-297) of Cue3 that forms a helical domain structure as predicted by AlphaFold2 (Fig. 3E, Suppl. Fig. 8B, Suppl. Fig. 9B and 9C). This domain of Cue3 is similar to the x-ray structure of the human homolog ASCC2 in complex with the N-terminal domain of ASCC3 (Slh1) which we do not observe (*28*). The predicted ubiquitin-binding CUE domain (298-360), a linked four-helix bundle (468-541), and the ultimate C-terminus (542-624) are not visible in our structures most likely due to their high flexibility. On the opposite convex side of Slh1, and in close vicinity to uS10, density for Rqt4 is visible (Fig. 2, Suppl. Fig. 9B). This density matches well with the predicted C2HC5-type zinc-finger domain (ZFD; 172-218) packing against the CTC RecA1 domain (Fig. 3B, 3F, Suppl. Fig. 8C, 9B and 9C). From there, additional, partially helical density extends towards the Slh1 cassette interface bridging the NTC WH domain to the CTC RecA2 domain. Here, the density matched well with the model for a region bridging the C-terminal part of the helical linker with the first helix of the winged-helix-like domain (Suppl. Fig. 8C and 9B). The remaining parts of Rqt4 including the N-terminal PWI-like domain (1-74), two α-helices flanking the ZFD as well as the C-terminus were not visible in our reconstruction due to flexibility.

Taken together, the RQT complex engages the 40S subunit of the ribosome in at least two different conformations, a main conformation C1 and a second conformation C2. In both conformations, RQT adopts an elongated architecture, stabilized by Cue3 and Rqt4 (Suppl. Fig. 10H) with the NTC of Slh1 providing the main anchor point on the ribosome involving mainly uS3 near rRNA h16 in both the C1 and C2 states. Meanwhile, the two accessory subunits Cue3 and Rqt4 bind Slh1 on opposite sides to bridge the NTC and CTC lobes. As a result, the RQT complex is positioned in immediate vicinity to the mRNA entry channel with the RNA substrate paths of its two helicase domains freely accessible to engage mRNA.

### RQT stabilizes an unusual 40S conformation

In the C2 state, the 80S ribosome adopted a conformation very similar to a (tRNA) translocation intermediate (TI) as described in a recent study analyzing eEF2-bound POST-states (*29*). The position of the 40S body relative to the 60S is similar to C1 and thus, closely resembles the classical POST-state. Yet, the 40S head in C2 has undergone a large counterclockwise swivel motion of 20 degrees, resembling essentially the TI-POST-2 state stabilized by eEF2 (*29*) (Fig. 4A-4C). While the previously observed translocation intermediate contains two tRNAs in chimeric ap/P and pe/E hybrid states, the RQT-bound C2 state ribosome has only one tRNA in the chimeric pe/E hybrid state as verified by signature interactions of the anticodon-stem loop with U1191 (*H.s*. U1248) and G904 (*H.s*. G961) (Fig. 4C).

The RQT in the head-swiveled TI-POST2 state (C2) showed the same inter-cassette arrangement as in C1. We interpret this to mean that RQT accompanies the swivel movement (rotation) of the 40S head essentially as a rigid body, but undergoes a small additional rotation towards the beak of the 40S head by about 10 degrees around its binding site on uS3 (Fig. 4A, 4B). As a consequence, whereas the main contacts to the 40S head via uS3 and the Slh1 FN3/HhH remain fully intact, the Slh1 NTC relocates on the body of the 40S subunit, with the trajectory of the 40S head swivel corresponding to a downwards movement of the NTC RecA domains of Slh1 along rRNA h16 (Fig.4B). The RecA2 domain thereby translocates from near the tip of h16 towards its stem.

In this new position, an additional new contact with h16 is formed by the FN3 domain that basically occupies the position of the HhH domain on h16 in the C1 state. In turn, the HhH forms a new rRNA contact with h18 at a position that is occupied by the C-terminal tail of eS30 in the C1 state (Suppl. Fig. 9E, 9F). Taken together, the Slh1 NTC is able to accommodate and stabilize two defined states on the 40S subunit with a distinct head-to-body arrangement.

### Application of force on mRNA corresponds to head-swiveling

Our structural observations show that RQT can engage 80S ribosomes in two conformations. Whereas one conformation represented a classical POST state 80S (C1), the second conformation (C2) represented an unusual late mRNA/tRNA translocation intermediate that so far has only been observed in presence of eEF2 trapped with non-hydrolyzable GMP-PNP and two tRNAs in ap/P and pe/E states (Fig. 4C) (*29*). During normal translocation, the step leading from the head-swiveled TI-POST states to the classical POST state is assumed to be the energy consuming step, where GTP-hydrolysis in eEF2 occurs (*29, 30*). The transition from our observed RQT-bound states C1 (POST) to C2 (head-swiveled TI-POST2) thus closely resembles a reverse translocation motion of the 40S head. We speculate that this motion of the 40S head could be key to the RQT-dependent splitting mechanism. Notably, C2 states were neither present in splitting incompetent reactions including an ATPase deficient Walker A mutation in the NTC of Slh1(K316R-RQT) nor in reactions with disomes lacking the 3’-mRNA overhang. From this we conclude that an ATP-dependent helicase activity on the mRNA must occur to acquire the C2 state, and that transition to this state is important for splitting of the lead ribosome of the stalled disome.

Interestingly, the transition from the POST-state lead ribosome into a head-swiveled C2 state could be easily achieved by applying force to the mRNA in the opposite direction of mRNA translocation (3’-to 5’-direction) – a pulling mechanism (*31*). This notion is also in agreement with the finding that in human cells mRNA damage results in collisions which are resistant to splitting by hRQT (*32*). Yet, the question remains whether both or only one of the two connected Slh1 cassettes are responsible for mRNA binding and pulling; based on sequence alignments, contrary to Brr2 (*33, 34*), Slh1 may have two active ATPase/helicase sites (Suppl. Fig. 9A), although DNA unwinding assays showed activity only for the NTC in human RQT (*28*). Since we found that single Walker mutations in both the NTC and the CTC abolish RQT activity (Fig. 1E), either both cassettes need to function as helicases, or one of the helicase activities depends on the other cassette’s ATPase cycle. Unfortunately, when analyzing the ATPase cassettes for presence of mRNA, no clear density could be identified in either one of them. Yet, when taking into account the relative position of the two cassettes with respect to the mRNA path at the 40S entry site, we speculate that only the CTC is fully capable of acting on emerging mRNA for several reasons: contrary to the CTC and also to Ski2 in the ribosome-bound SKI complex (*22, 23*) the NTC is positioned on the ribosome such that its RNA entry channel is facing away from the 40S mRNA entry site (Suppl. Fig. 10A-D); this becomes clear when comparing ribosome-bound Slh1 to RNA-bound Brr2 (*35*) (Suppl. Fig. 10G). Assuming the same directionality of the helicases and a direct threading of mRNA into the helicase core, the Slh1 NTC activity would result in pushing the mRNA in the 5’-3’ direction rather than applying a pulling force. To realize a pulling (or extraction) activity, the mRNA would need to wind along the inner side of h16 and around the RecA2 domain to enter the helicase core from the other side; this seems sterically highly unlikely. Moreover, in the C1 conformation, direct entry for the mRNA into the NTC helicase core, as observed in the Ski-complex, would be obstructed by rRNA h16 that resides at the interface of RecA1 and RecA2, occupying the space where RNA would exit the helicase (Fig. 4D and Suppl. Fig. 10D, E). In addition, in the C1 conformation, the helicase core of the NTC is in a constricted conformation, mostly resembling the apo/unengaged states of Brr2 and other related Ski2-like helicases. Therefore, the position and conformation of the Slh1 NTC on the 40S probably prevents engagement with mRNA rather than facilitating it. We speculate that the NTC ATPase activity may be required for the transition on the rRNA h16.

We propose that the CTC represents the helicase unit responsible for engaging and applying pulling force on the mRNA. The CTC is positioned ideally to engage with the substrate mRNA in a way such that pulling would occur; and, when superimposing the mRNA path as visible in the ribosome-SKI complex onto RQT-C2, the mRNA would indeed directly thread into the CTC helicase channel (Suppl. Fig. 10E). Moreover, a conformational transition from C1 to C2 corresponds to a shortening of the distance from the CTC helicase unit of Slh1 to the ribosomal mRNA entry by about 7 Å (Fig. 4E). The application of a pulling force to the mRNA by the CTC helicase may be sufficient to trigger the substantial conformational transition from C1 to C2. Importantly, however, we can neither provide direct evidence for the CTC acting as the mRNA helicase nor can we exclude a helicase function of the NTC. The exact roles of the individual ATPase cassettes will require further clarification.

### Mechanistic model for RQT-dependent ribosome splitting

Together with previous findings, our observations allow us to propose an initial working model for the molecular mechanism underlying RQT-dependent splitting of the lead ribosome (Fig. 5). In the first step, the RQT is recruited to the ribosome queue after Hel2-dependent K63-linked uS10 polyubiquitination, likely via interaction with the CUE domain of the Cue3 subunit (*7, 36, 37*). After recruitment, RQT stably locks onto the 40S of the stalled/lead ribosome in the C1 conformation via the Slh1 NTC, placing the Slh1 CTC in a position primed to bind the 3’ region of the mRNA emerging from the ribosomal mRNA entry site. The ATPase dependent pulling force and (partial) extraction of the mRNA by the Slh1-CTC (and/or NTC) then causes a head-swivel movement on the lead ribosome and the relocation of Slh1’s NTC. This head swiveling results in re-positioning the P/P site tRNA into an ap/P conformation (and the E/E site tRNA in a pe/E conformation), which likely renders the POST-TI2 80S substantially less stable than the POST-state 80S; this idea is supported by the observation that this intermediate has only been observed with eEF2 trapped in the presence of non-hydrolyzable GMP-PNP. Such a state may lead to a destabilized 80S primed for dissociation of the 40S from the peptidyl-tRNA carrying 60S. Yet, this is not sufficient since we see that for efficient splitting, the presence of a trailing ribosome is necessary.

In our study, we observe that head-swiveling directly exerts force on the trailing collided ribosome as seen in the induced disorder of the RACK1/Asc1-RACK1/Asc1 contact (Fig. S7D, E; Suppl Movie). In a trajectory of the swiveling motion, RACK1/Asc1 of the lead ribosome would clash with RACK1/Asc1 of the trailing ribosome, forcing the trailing ribosome to accommodate this movement by rotating in the opposite direction with the connecting mRNA between the two ribosomes serving as a pivot. This results in a net movement of the trailing ribosome in the direction of the 60S subunit of the lead ribosome, with a primary impact on the contact of the 40S-body to the stalled 60S of the disome. In this situation, we speculate that the 60S of the lead ribosome will be dislocated from its position, leading to further destabilization of the lead ribosome and eventual dissociation of the 60S subunit from the 40S. Moreover, in case of prolonged RQT helicase activity and extended pulling on the mRNA, the trailing ribosome would ultimately serve as a giant wedge that is driven between the subunits of the lead ribosome. Yet, since the mRNA connecting the collided ribosomes must be already under tension as indicated by the incomplete tRNA translocation of the trailing ribosomes, we predict that any additional force application, extending the mRNA by only a few nucleotides, should have an immediate destabilizing effect.

The proposed RQT mechanism is reminiscent of the remodeling of the spliceosome by the two related RNA helicases Brr2 and Prp22. Brr2 engages and translocates a single-stranded region of U4 snRNA (*38*), and Prp22 applies pulling force on single stranded mRNA for its release (*31*). However, a ribosome splitting mechanism that is driven by an mRNA helicase and that works through destabilizing conformational transitions of the ribosome, represents a novel principle. For ribosome recycling after canonical translation termination, bacteria employ the GTPase EF-G together with the cofactor RRF (ribosome recycling factor), which functions by driving the cofactor as a splitting wedge between the ribosomal subunits (*39-41*). The ABC-type ATPase ABCE1 works in a similar way in *Archaea* and eukaryotes, also by causing steric clashes between the subunits and cofactors aRF1, eRF1 and Dom34, respectively (*42, 43*). However, the ABCE1 machinery cannot function on stalled collided ribosomes as long as a tRNA in the A site or a translation factor prevent cofactor binding. This may explain the necessity for the evolution of an alternative splitting mechanism employing a mechanism that can function on any stalled ribosome irrespective of its A site occupancy. Importantly, the observed mechanism is highly specific in that it works only on ribosomes with a collided neighbor and requires licensing/recruitment through ubiquitin tagging of the stalled ribosome. These ideas are in line with the recent discovery in the bacterium *Bacillus subtilis* of an alternative splitting factor, the ATPase MutS2, that clears collided ribosomes (*44*). Although its mode of function is not entirely clear yet, it appears to function independent of mRNA, thereby making the RQT mechanism a unique example of an mRNA helicase dependent ribosomal splitting device which is conserved in eukaryotes from fungus to humans. Further studies on RQT will shed light on the exact mode of mRNA engagement and on the role of the ATP hydrolysis cycle of the individual helicase units in promoting ribosome splitting.

## Materials and Methods

### RQT purification

A yeast strain overexpressing RQT components (Slh1-FTP, Cue3, Rqt4, see Table S2) was grown in synthetic drop-out media. Cells were harvested by centrifugation and subsequently disrupted by cryogenic milling. Cell powder was resuspended in lysis buffer (50 mM K_2_HPO_4_ /KH_2_PO_4_ pH 7.5, 500 mM NaCl, 5 mM Mg(OAc)_2_, 100 mM arginine, 1 mM DTT, 1 mM PMSF, 0.1 % NP-40, 1 protease inhibitor pill/50 ml (Roche, #04693132001)) and centrifuged at 30,596 x *g* for 30 min. Purification was performed on IgG-sepharose (GE Healthcare, #GE17096901) resin. The complex was eluted by TEV cleavage in RQT elution buffer (50 mM HEPES pH 7.5, 300 mM KOAc, 5 mM Mg(OAc)_2_, 0.01% Nikkol, 1 mM DTT) for 1.5 h at 4 °C.

### eIF6 purification

*S. cerevisiae* eIF6A was purified essentially as described before (*45*) from the p7XC3GH (Addgene #47066; Watertown, MA, USA) plasmid, expressing eIF6 with a C-terminal 3C protease cleavage site, GFP, and His_10_-tag from *E. coli* Rosetta cells.

### Ubc4 purification

Recombinant Ubc4 was purified as GST-Ubc4 fusion protein from *E. coli* Rosetta-gami 2 (DE3) cells harbouring the pGEX-Ubc4 plasmid essentially as described before (*46*).

### Hel2 purification

A yeast strain (see Table S2) overexpressing Hel2-FLAG was grown in synthetic dropout media. Cells were harvested by centrifugation and subsequently disrupted by cryogenic milling (SPEX SamplePrep 6970EFM Freezer/Mill). Cell-powder was resuspended in lysis buffer (50 mM Tris pH 7.5, 500 mM NaCl, 10 mM Mg(OAc)_2_, 0.01 % NP-40, 1 mM PMSF/DTT, 1 protease inhibitor pill/50 ml (Roche, #04693132001)) and centrifuged at 30,596 x *g* for 30 min. The supernatant was purified using M2 FLAG affinity resin (Sigma Aldrich, #A2220). During the washing steps, the salt concentration was decreased from 500 mM NaCl to 100 mM NaCl in 100 mM steps. Hel2-FLAG was eluted by incubation of the resin with 3x FLAG peptide in elution buffer (50 mM HEPES, 100 mM KOAc, 5 mM Mg(OAc)_2_, 1 mM DTT, 0.05% Nikkol).

### *In vitro* translation of stalling constructs

CGN_12_-CGN_6_- and CGN_4_-reporters were generated by PCR using the plasmid pEX-His-V5-uL4a-CGN12 as template. It expresses a truncated version of ribosomal protein uL4 (*47*) followed by a mRNA-based stalling sequence containing twelve consecutive arginine-encoding CGN (N=A, G or C) codons (CGN_12;_ “R12 cassette”) and a 3’-region of 129 bases. CGN_6_ and CGN_4_ constructs were derived by truncation after the sixth or fourth CGN codon. *In vitro* translation of these constructs would result in accumulation ribosomes stalled after the second or third CGN codon in the P-site (*7*) with long (CGN_12_) or short (CGN_6_; CGN_12_) 3’-mRNA overhangs emerging from the mRNA entry channel of the stalled ribosome (see Fig. 1 and Fig. S1).

*His-V5-uL4-CGN* mRNAs were obtained using a mMESSAGE mMACHINE T7 kit (Invitrogen #AM1344) using linear DNA templates. Cell-free yeast translation extracts were prepared as described previously using Δ*xrn1*Δ*slh1*Δ*cue2* or Δ*ski2*/*uS10-HA* yeast strain (*48, 49*). The *in vitro* translation reaction was performed in the presence of Hel2 (50 nM) at 17 °C for 75 min. Ribosome nascent chain complexes (RNCs) were purified via the encoded His-Tag using magnetic Dynabeads (Invitrogen, #10104D). The translation reaction was applied to equilibrated beads (50 mM HEPES pH 7.5, 300 mM KOAc, 10 mM Mg(OAc)_2,_ 125 mM sucrose, 0.01 % NP-40, 5 mM β-mercaptoethanol) for 20 min, washed and eluted in elution buffer (50 mM HEPES pH 7.5, 150 mM KOAc, 10 mM Mg(OAc)_2_, 125 mM sucrose, 0.01 % NP-40, 5 mM β-mercaptoethanol) containing 400 mM imidazole.

### Isolation of di- and trisomes using sucrose density gradients

Purified RNCs were loaded on top of 10-50 % sucrose gradients (50 mM HEPES pH 7.5, 200 mM KOAc, 10 mM (MgOAc)_2_, 1 mM DTT, 10/50 % sucrose(w/v)) followed by ultracentrifugation for 3 h at 284,000 x *g*. Fractions of the gradients were collected on a gradient station (Biocomp) equipped with a TRIAX flow cell (Biocomp) and a GILSON fractionator. Mono-, di- and trisome fractions were used for further experiments.

### *In vitro* ubiquitination of stalled RNCs

The *in vitro* ubiquitination reaction was performed essentially as described before with adjustments(*46*). 10-15 pmol of purified RNCs, or isolated di-/trisomes were incubated with 57 µM ubiquitin, 116 nM UBE1 (R&D systems), 3.3 µM Ubc4 and 757 nM Hel2 in a reaction buffer (20 mM HEPES-KOH pH 7.4, 100 mM KOAc, 5 mM (MgOAc)_2_, 1 mM DTT, 1 mM ATP, 10 mM creatine phosphate, 20 ug/ml creatine kinase (Roche)) at 25 °C for 30 min. Ubiquitinated ribosomes were either used directly in *in vitro* splitting assays or pelletted through a sucrose cushion (50 mM HEPES pH 7.5, 100 mM KOAc, 25 mM Mg(OAc)_2_, 1 M sucrose, 0.1 % Nikkol) in a TLA110 rotor (Beckman Coulter) at 434,513 x *g* for 1.5 h at 4 °C, for the subsequent preparation of cryo-EM samples.

### Semi-dry Western blotting

After sample separation on NuPAGE, semi dry western-blotting was performed. The PVDF membrane was blocked with 5% skim milk in TBS for 1 h and incubated with anti-HA-HRP (Roche, 3F10, 1:5000) in 5% milk/TBS. After washing (1x TBS with 0.1 % (w/v) TWEEN-20, 2x TBS) signal was detected with an AI-600 imager (GE Healthcare) using SuperSignal West Dura Extended Duration Substrate (Thermo).

### *In vitro* splitting assays

6-12 pmol of RNCs (ubiquitinated or non-ubiquitinated) were incubated with 10x molar excess of RQT complex (wild type or K316R-Slh1 mutant; RQT*) and 5x molar excess of eIF6 for 15 min at 25 °C. For control reactions without RQT, RQT elution buffer (50 mM HEPES pH 7.5, 300 mM KOAc, 5 mM Mg(OAc)_2_, 0.01% Nikkol, 1 mM DTT) was added instead. The reactions were separated on a 10 – 50 % sucrose gradient using ultracentrifugation in a SW40 rotor for 3 h at 284,000 x *g*.

### CGN_12_-reporter gene assay to probe for CAT-tailin

Exponentially grown yeast cultures were harvested at OD_600_ of 0.5 ∼ 0.8. Cell pellets were resuspended with ice-cold TCA buffer (20 mM Tris pH8.0, 50 mM NH_4_OAc, 2 mM EDTA, 1 mM PMSF, 10% TCA) and then an equal volume of 0.5 mm dia. zirconia/silica beads (BioSpec) was added followed by thorough vortexing for 30 s, three times at 4 ºC. The supernatant was collected in a new tube. After centrifugation at 18,000 x *g* at 4 ºC for 10 min and removing supernatant completely, the pellet was resuspended in TCA sample buffer (120 mM Tris, 3.5% SDS, 14% glycerol, 8 mM EDTA, 120 mM DTT and 0.01% BPB). Proteins were separated by SDS-PAGE and were transferred to PVDF membranes (Millipore; IPVH00010). After blocking with 5 % skim milk, the blots were incubated with the anti-GFP antibody (sc9996, Santa Cruz), followed by the 2^nd^ incubation with the anti-mouse IgG antibodies conjugated with horseradish peroxidase (NA931V, Cytiva). The products derived from CGN_12_ reporter gene were detected by homemade ECL solution using the ImageQuant LAS4000 (GE Healthcare).

### Cryo-EM samples and data collection of RQT-ribosome complexes

To generate suitable samples for cryo-EM, at least 6 pmol of ubiquitinated disomes were incubated for 5 min at 25 °C with 12 pmol of purified RQT complex in the presence of 1 mM ATP. After incubation, 3.5 µl of samples were vitrified in liquid ethane on glow discharged, R3/3 copper grids with a 2 nm carbon coating (Quantifoil) using a Vitrobot mark IV (FEI) with 45 s wait time and 2.5 s blotting time.

Altogether four samples were analyzed, two pre-splitting reactions and two post-splitting reactions. Pre-splitting reactions contained either ubiquitinated CGN_6_-stalled disomes (with short accessible 3’-mRNA) and wild type RQT (RQT_wt_) or (ubiquitinated) CGN_12_-stalled disomes (with long accessible 3’-mRNA) and the RQT mutant containing K316R-Slh1 (RQT*) as well as eIF6. Post-splitting reactions contained ubiquitinated CGN_12_-stalled disomes and wild type RQT, one reaction with and one without eIF6.

For the post-splitting sample (with CGN_12_-stalled disomes and RQT_wt_ without eIF6 addition) sample, 21.171 movies were collected on a Titan Krios with a K2 Summit DED, at 300 keV with a pixel size of 1.045 Å/pixel. The applied electron dose was approximately 1.09 e^-^/Å/frame for 40 frames and data were collected in a defocus range between 0.5-3.5 µm. All frames were gain corrected, aligned and subsequently summed using MotionCor2 (*50, 51*). Downstream data processing was performed using CryoSPARC (v.3.3.1) (*52, 53*). The CTF was estimated using gCTF and CTFFIND4 (*54, 55*), followed by particle picking via CryoSPARCs blob picker. After 2D classification, 2.415.630 particles were used for 3D refinement. Subsequent rounds of classification were carried out using 3D Variability Analysis (*53*) with a soft mask around the ribosomal 40s subunit. Subsequent sorting steps with a soft mask around the RQT complex yielded two classes containing RQT in two distinct conformations (C1 and C2). The class of 80S ribosomes with RQT bound in C1 contained 194.186 particles (8 % of total particles) and was refined to a overall resolution of 2.4 Å. Subtracted particles of RQT were locally refined to a resolution of 3.5 Å. The second class of 80S with RQT bound in C2 contained 20.380 particles and was refined to a overall resolution 3.0 Å. Local refinement was carried out as for C1 and resulted in a resolution of 4.8 Å. The processing scheme and (local) resolution estimations for sample are depicted in Fig. S4 and Fig. S6, respectively.

The other three samples were vitrified and and cryo-EM data were obtained as described above. For the CGN_12_/RQT* pre-splitting sample, 11.251 movies were collected with an applied electron dose of approximately 1.09 e^-^/Å/frame for 40 frames. For CGN_6_/RQT_wt_/eIF6 pre-splitting sample, 16.508 movies were collected with an applied electron dose of approximately 1.1 e^-^/Å/frame for 40 frames. For CGN_12_/RQT_wt_/eIF6 post-splitting sample, 14.092 movies were collected with an applied electron dose of approximately 1.16 e^-^/Å/frame for 40 frames. Downstream processing was performed using CryoSPARC (v3.3.1) as depicted in Figs S2-S5.

### Model building and refinement

The RQT-ribosome C1 model was prepared by rigid body docking the model for *SDD1*-stalled 80S (PDB code 6SNT; (*8*)) and the models predicted by Alpafold 2 (*24, 56*) for RQT components Slh1 (Uniprot-ID P53327), Cue3 (Uniprot-ID P53137) and Rqt4 (Uniprot-ID P36119). For Slh1, the N-terminal and C-terminal cassettes (NTC; residues 217-1122 and CTC residues 1123-1967) were docked individually. For Cue3 only the N-terminal part (residues 1-297) and for Rqt4, two parts (residues 171-219 and 323-381) were docked (see also Fig. S8).

To obtain the RQT-ribosome C2 model, 60S, 40S and RQT of the RQT-ribosome C1 model were individually rigid-body docked into the cryo-EM density. For the collided ribosomes, 40S and 60S of the *SDD1* stalled 80S model (PDB code 6SNT) were docked individually. tRNAs were taken from the model of the yeast disome (PDB code 6I7O) (*46*) and rigid body docked into the A/P, P/E site densities.

The 60S model was prepared by docking the model for NatA-bound 60S (PDB code 6HD7) (*47*) into the cryo-EM density. tRNA was taken from 6SNT, eIF6 from PDB code 1G62 (*57*).

Adjustment of all models was performed using Wincoot (v.0.9.6) and subsequently real-space refined using Phenix (1.19.1) (*58*). To obtain figures, molecular models and cryo-EM densities were displayed in ChimeraX (v.1.3) (*59, 60*)

## Acknowledgments

We thank C. Ungewickell, S. Rieder and Alicia Musial for excellent technical assistance, Dr. Rachel Green for kindly providing the Δ*cue2*, Δ*slh1*, Δ*xrn1* yeast knockout strain. Dr. Petr Tesina and Timo Denk for fruitful discussions and Dr. Jingdong Cheng for advice in model building.

## Author contributions

R.B., T.I., K.B., and T.B. designed the study; O.B. collected cryo-EM data; K.B. and L.K. prepared cryo-EM samples and processed cryo-EM data; K.B. built molecular models; Y.M. performed the CGN12 reporter gene assay. K.B., L.K. and D.B. performed in vitro translation and splitting assays with help from J.M. and Y.M.; K.B. and K.I. performed *in vitro* ubiquitination. K.I. and Y.M. generated plasmids and yeast strains. K.I., K.B. D.B. and J.M. performed protein purifications; T.B., R.B. T.I. and K.B. wrote the manuscript, with comments from all authors.

## Data and materials availability

Cryo-EM maps and molecular models have been deposited at the Electron Microscopy Data Bank and Protein Data Bank with accession codes EMD-XXXX and PDB-YYYY. Materials are available from the authors on request.

## Supplementary Materials

Figs. S1 to S10

Tables S1 to S2

Movie S1

