## Supplementary Material - Figures and Tables for "Clearing of ribosome collisions by the ribosome quality control trigger complex RQT"

**This PDF file includes:**

Figs. S1 to S10
Tables S1 to S2
Captions for Movie S1

**Other Supplementary Materials for this manuscript include the following:**

Movies S1

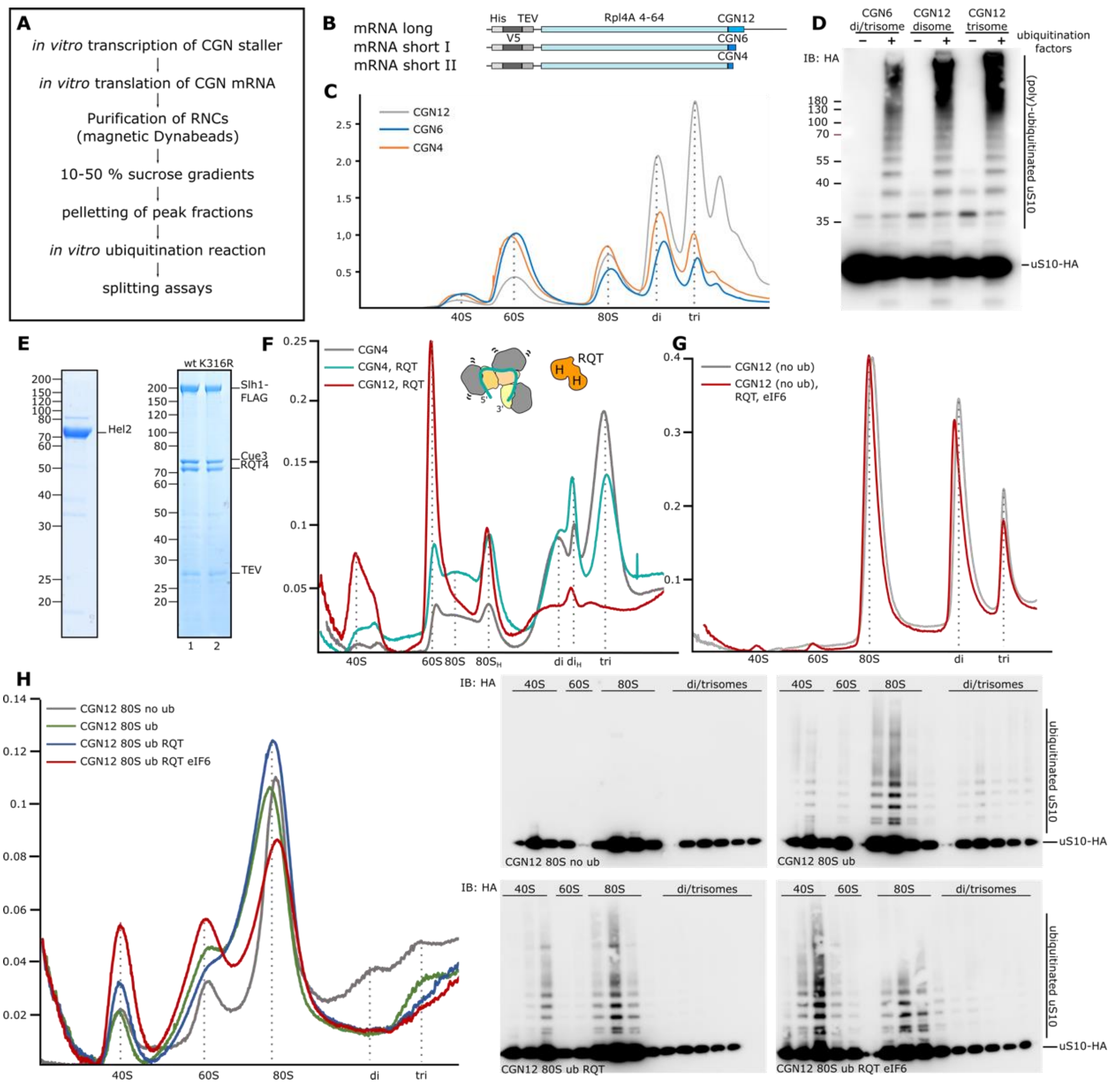


**Fig. S1: Experimental setup and controls of the RQT-dependent in vitro splitting assay.**

(**A**) Experimental outline for generation of ubiquitinated di- and trisomes to be used in splitting assays. (**B**) Schematic representation of (CGN)-stalling mRNA constructs. The $CGN_{12}$ stalling sequence is followed by a stop codon to release ribosomes reading the stall sequence and then followed 138 nucleotides of 3'-UTR. This results in 147 bases downstream of the stall site (the second or third CGN codon in the ribosomal A-site). $CGN_6$ and $CGN_4$ constructs are truncated versions of CGN12 with 9-12 and 3-6 nt downstream the stall site. (**C**) Polyribosome profiles of CGN-stalled RNCs after sucrose density gradient centrifugation. Di-and trisome peaks were used for in vitro ubiquitination (**D**) Western Blot analysis of CGN-stalled trisomes obtained from a yeast strain harboring HA-tagged uS10 ribosomes before and after in vitro ubiquitination. (**E**) SDS-PAGE gel showing purified Hel2 and RQT complex (wt = wild type; RQC* = K316R mutant). (**F**) *In vitro* splitting assays comparing $CGN_4$-stalled trisomes with no 3'-mRNA emerging from the mRNA entry of the stalled ribosome with $CGN_{12}$-trisomes (long 3'-mRNA stretch). (**G**) In vitro splitting assay using $CGN_{12}$-trisomes (long 3'-mRNA stretch) before in vitro ubiquitination. (**H**) *In vitro* splitting assay comparing ubiquitinated and non-ubiquitinated $CGN_{12}$-stalled 80S-RNCs (harboring HA-tagged uS10) in presence and absence of eIF6. Left: polyribosome profiles; right anti-HA Western Blots from gradient fractions. Note that these RNCs mainly consist of 80S ribosomes and that substantial amounts of the ubiquitinated 80S are not split by RQT.

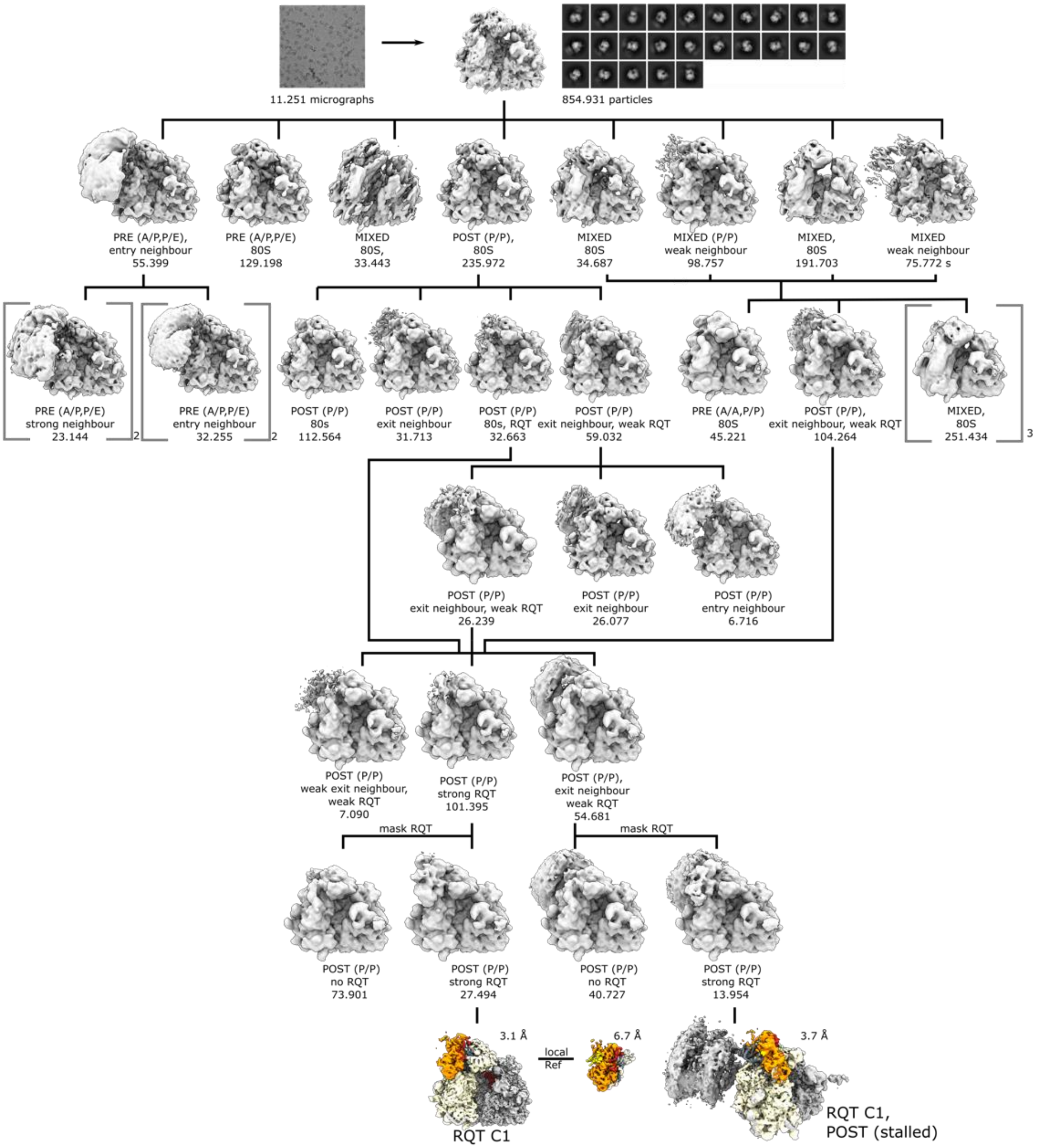


**Fig. S2: Sorting scheme for cryo-EM analysis of the pre-splitting reaction using K316R-RQT and disomes with a long 3' mRNA stretch ($CGN_{12}$) in presence of eIF6.**

From a total of 11.251 micrographs, approx. 855.000 ribosomal particles were selected after 2D classification and a consensus refinement was performed. This was followed by exhaustive heterogeneity assessment using the CryoSPARC-2 3D variability analysis tool. The outcome is summarized here. Eight ribosomal classes were obtained and the most prominent classes represented ribosomes in the non-rotated post-translocational (POST) state with a P/P site tRNA and rotated pre-translocational state (PRE) with hybrid A/P-P/E tRNAs. Individual classes also differed in presence or absence of a neighboring ribosome, either on the mRNA entry site (mostly for hybrid state 80S) or the mRNA exit site (mostly for POST state 80S), and in presence or absence of density for RQT on the 40S head. Several classes still represented mixed states yielding in lower resolution reconstructions. Similar classes were joined and further sub-classified. This resulted in three stable classes of POST-state ribosomes with extra density for RQT. Focused 3D variability analysis using a soft mask covering the head region of the ribosome yielded two homogenous classes, one displaying a strong neighbor at the mRNA exit and one 80S monosome. The 80S was refined to high resolution followed by local refinement of the RQT density yielding in a final resolution of 3.1 Å for the 80S and 6.7 Å for the RQT complex. RQT is bound to the lead ribosome in the "C1" state. By shifting the center of the extracted box, and subsequent refinement, also the disome was reconstructed from the strong neighbor class, resulting in a disome with RQT bound to the lead ribosome in the C1 state. Due to limited amount of particles, the disome was not further refined.

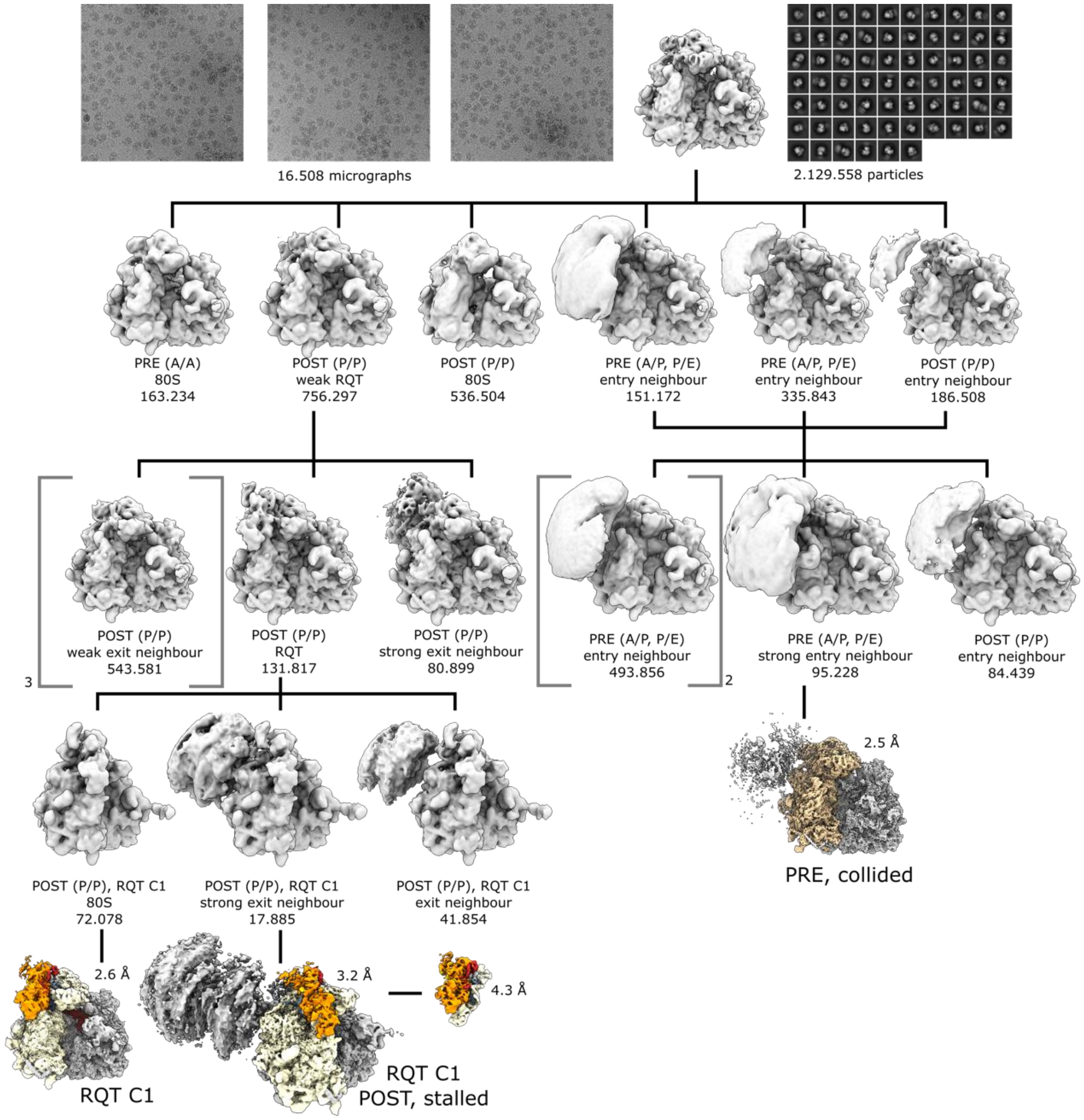


**Fig. S3: Sorting scheme for cryo-EM analysis of the pre-splitting reaction using disomes with a short 3’ mRNA stretch ($CGN_6$) in presence of eIF6.**

From a total of 16.508 micrographs, 2.13 million ribosomal particles were selected after 2D classification and a consensus refinement was performed. This was followed by exhaustive heterogeneity assessment using the CryoSPARC-2 3D variability analysis tool. Six main ribosomal classes were distinguished, mostly representing ribosomes in the non-rotated post-translocational (POST) state with a P/P site tRNA and rotated pre-translocational state (PRE) with hybrid A/P-P/E tRNAs. Individual classes also differed in presence or absence of a neighboring ribosome, either on the mRNA entry site (mostly for hybrid state 80S) or the mRNA exit site (mostly for POST state 80S). One class of the POST-state 80S showed additional density on the 40S for RQT. This large class (756.297 particles) was further sub-sorted resulting in three stable RQT containing classes, one with a strong neighbor at the mRNA exit, one with a weak neighbor in the same position and one 80S monosome. By shifting the center of the extracted box, and subsequent refinement, a disome was reconstructed from the strong neighbor class. The lead ribosome was refined to high resolution followed by local refinement of the RQT density yielding in a final resolution of 3.2 Å for the 80S and 4.3 Å for the RQT complex. RQT is bound to the lead ribosome in the “C1” state. From further sorting of PRE-ribosome classes, one class with a strong density for a neighbor on the mRNA entry site – representing a collided ribosome - was selected and refined to an overall resolution of 2.5 Å. This reconstruction was used to obtain a molecular model for the $(CGN)_{12}$ stalled disome (see Mat. & Meth.).

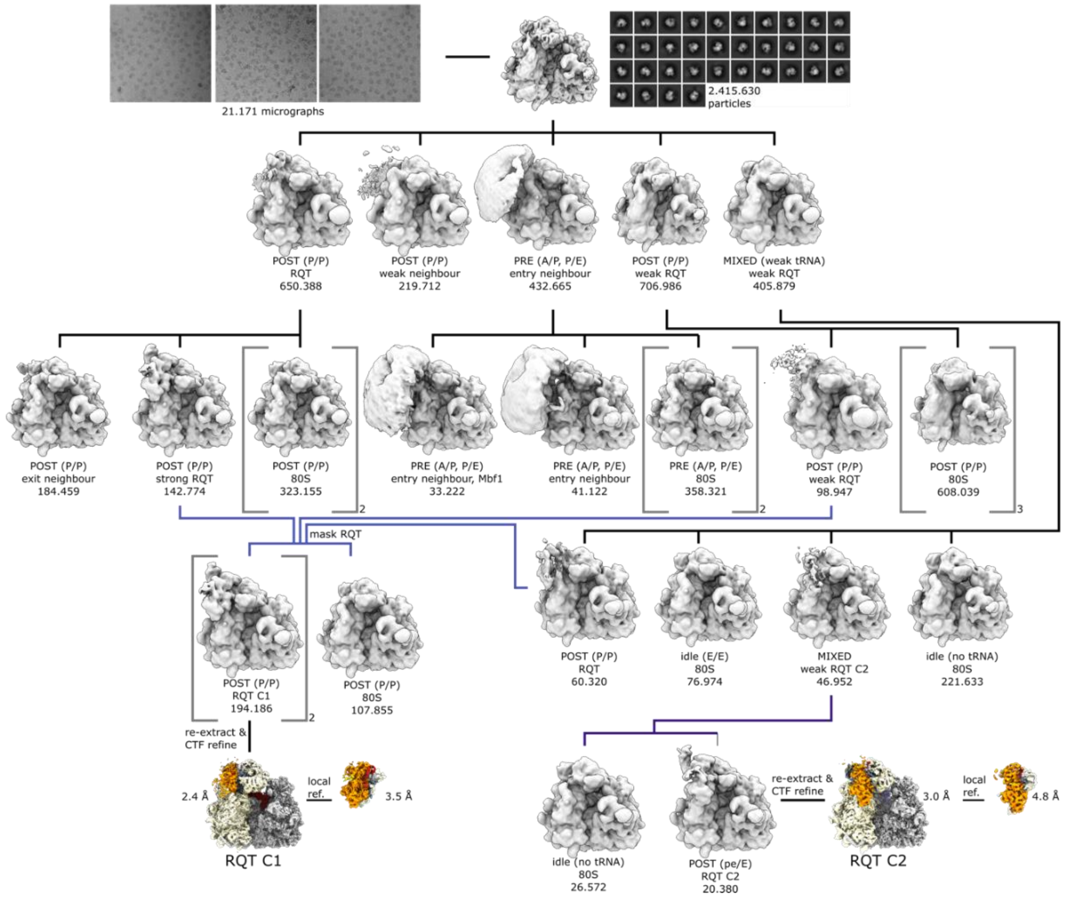


**Fig. S4: Sorting scheme for cryo-EM analysis of the post-splitting reaction using disomes with a long 3' mRNA stretch ($CGN_{12}$) in absence of eIF6.**

From a total of 21.171 micrographs, 2.41 million ribosomal particles were selected after 2D classification and a consensus refinement was performed. This was followed by exhaustive heterogeneity assessment using the CryoSPARC-2 3D variability analysis tool. Five main ribosomal classes were distinguished. They represented ribosomes in the non-rotated post-translocational (POST) state with a P/P site tRNA, rotated pre-translocational state (PRE) with hybrid A/P-P/E tRNAs, and mixed classes with low-tRNA occupancy and low-resolution. Individual classes also differed in presence or absence of a neighboring ribosome, either on the mRNA entry site (mostly for hybrid state 80S) or the mRNA exit site (mostly for POST state 80S) and extra density for RQT on the 40S head. Four of the five primary classes were sub-classified further. Resulting POST state ribosomes containing RQT were merged and sub-sorted using soft masks around the 40S head region, yielding two classes with highly enriched RQT density. These classes were merged and refined followed by local refinement for RQT after signal subtraction, yielding in a final resolution of 2.5 Å for the 80S (representing the C1 state) and 3.5 Å for the RQT complex. The RQT-containing 80S in the C2 state was obtained from the weak tRNA class, that contained idle ribosomes with only traces of tRNA which could be separated into RQT-containing particles in both C1 and C2 states. The C2 state particles were refined followed by local refinement of the RQT complex yielding in a final resolution of 3.0 Å for the 80S (representing the C2 state) and 5.0 Å for the RQT complex. Notably, despite some classes showing density for a neighboring ribosome, no stable disomes with RQT could be reconstructed from this dataset.

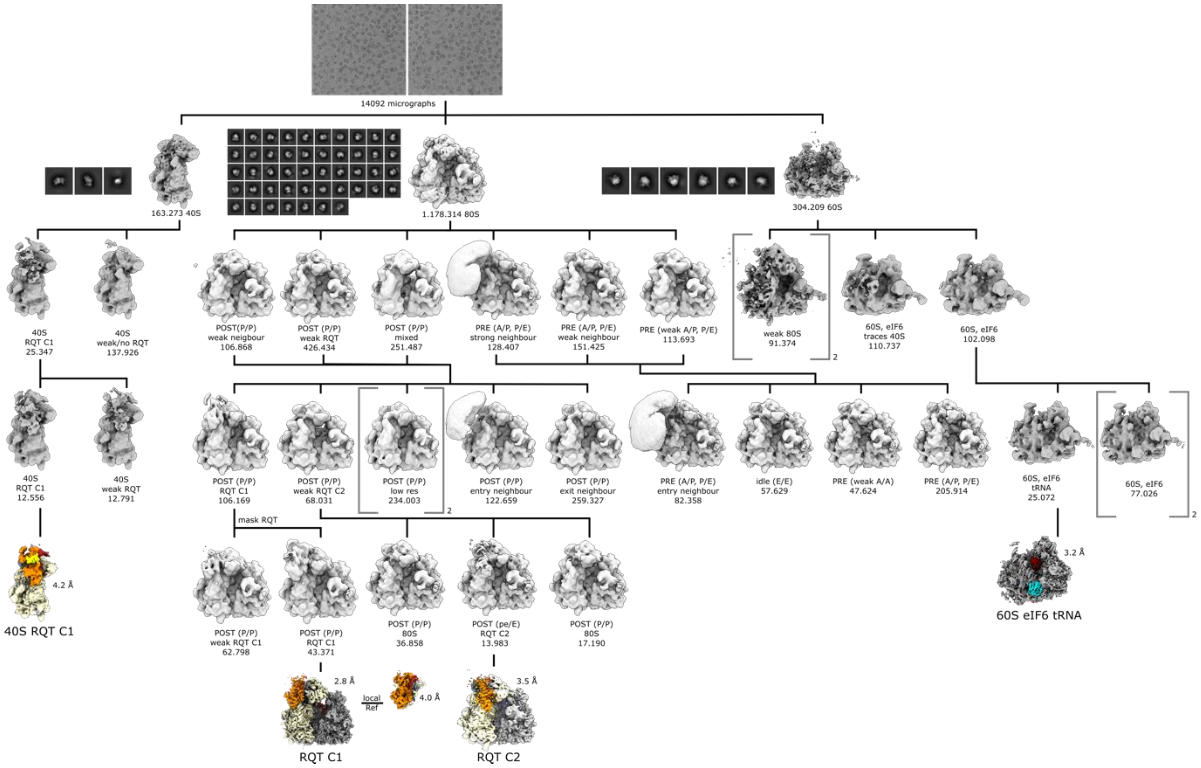


**Fig. S5: Sorting scheme for cryo-EM analysis of the post-splitting reaction using disomes with a long 3' mRNA stretch ($CGN_{12}$) in presence of eIF6.**

From a total of 14.092 micrographs, 1.18 million 80S, 304.209 60S and 163.273 40S ribosomal particles were selected after 2D classification and a consensus refinements were performed. This was followed by exhaustive heterogeneity assessment using the CryoSPARC-2 3D variability analysis tool. For the 80S, six ribosomal classes were initially distinguished. They represented ribosomes in the non-rotated post-translocational (POST) state with a P/P site tRNA, rotated pre-translocational state (PRE) with hybrid A/P-P/E tRNAs, and mixed classes with low-tRNA occupancy and/or low-resolution. Individual classes also differed in presence or absence of a neighboring ribosome, either on the mRNA entry site (mostly for hybrid state 80S) or the mRNA exit site (mostly for POST state 80S) and extra density for RQT on the 40S head. Similar classes were joined and sub-classified further. RQT-containing particles were locally classified using a soft mask on the 40S head region. This approach finally yielded in two RQT containing 80S classes, C1 and C2.  The C1 class was refined followed by local refinement of the RQT complex yielding in a final resolution of 2.8 Å for the 80S (representing the C2 state) and 4.1 Å for the RQT complex. The C2 class was refined to and overall resolution of 3.5 local refinement of RQT did not yield in improved resolution, likely due to limiting amount of particles. Sub-classification of the 40S particles resulted in one class with RQT bound in the C1 conformation. Sub-classification of the 60S particles yielded in a subset of eIF6-containing particles, a fraction of which contained density of tRNA bound in the P-site. This data subset was refined to 3.2 Å. Notably, despite some classes showing density for a neighboring ribosome, no stable disomes could be reconstructed from this dataset.

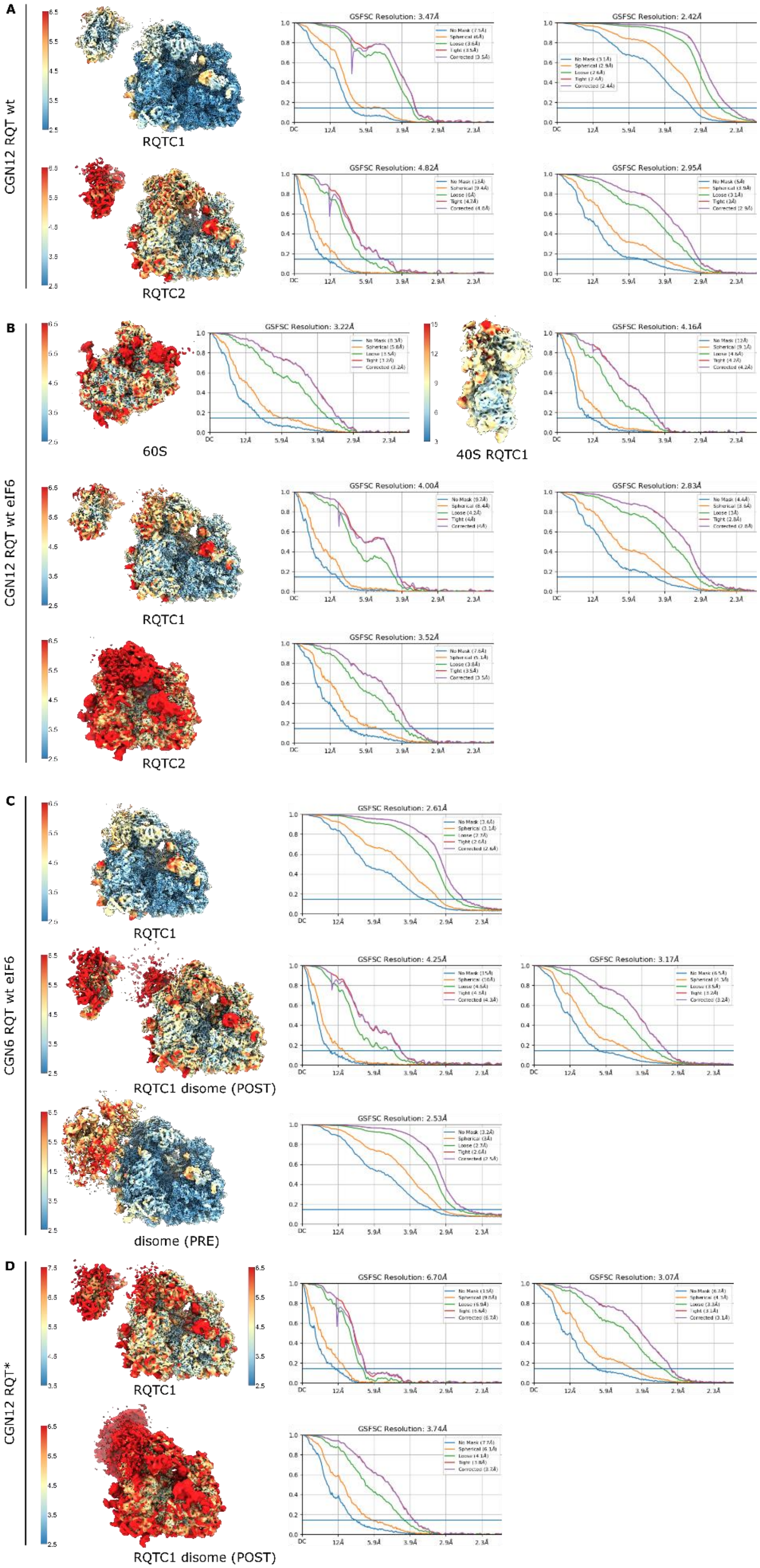
A
CGN12 RQT wt
RQTC1
GSFSC Resolution: 3.47Å
GSFSC Resolution: 2.42Å
RQTC2
GSFSC Resolution: 4.82Å
GSFSC Resolution: 2.95Å
B
CGN12 RQT wt eIF6
60S
GSFSC Resolution: 3.22Å
40S RQTC1
GSFSC Resolution: 4.16Å
RQTC1
GSFSC Resolution: 4.00Å
GSFSC Resolution: 2.83Å
RQTC2
GSFSC Resolution: 3.52Å
C
CGN6 RQT wt eIF6
RQTC1
GSFSC Resolution: 2.61Å
RQTC1 disome (POST)
GSFSC Resolution: 4.25Å
GSFSC Resolution: 3.17Å
disome (PRE)
GSFSC Resolution: 2.53Å
D
CGN12 RQT*
RQTC1
GSFSC Resolution: 6.70Å
GSFSC Resolution: 3.07Å
RQTC1 disome (POST)
GSFSC Resolution: 3.74Å

**Fig. S6: Overall and local resolution of the main classes present in the pre- and post-splitting reactions**

Cryo-EM maps and resolution curves are shown for all ribosomal particles relevant in the RQT-dependent splitting reaction. (**A**) shows RQT-80S-C1 and RQT-80S-C2 complex and the locally refined RQT complex as obtained from the post-splitting sample with ubiquitinated $CGN_{12}$-disomes and wt RQT in the absence of eIF6. (**B**) shows the RQT-40S, the eIF6-tRNA-60S and the two RQT-80S-complexes (C1 and C2) and respective locally refined RQT as obtained from the post-splitting sample with ubiquitinated $CGN_{12}$-disomes wt RQT in the presence of eIF6. (**C**) shows RQT-80S-C1 and the RQT-bound disome and respective locally refined RQT. The disome was locally refined on the RQT-bound lead (POST-state) 80S and the collided (PRE-state) ribosome. These maps were obtained from the pre-splitting sample with ubiquitinated $CGN_6$-disomes (short 3'-mRNA stretch) and wt RQT in the presence of eIF6. (**D**) shows RQT-80S-C1, locally refined 80S-bound RQT and the RQT-bound disome, refined on the lead (POST-state) ribosome. These maps were obtained from the pre-splitting sample with ubiquitinated $CGN_{12}$-disomes (long 3'-mRNA stretch) and ATPase deficient K316R-RQT (RQT*) in the presence of eIF6. All maps are filtered and colored according to local resolution. Resolution curves were exported from CryoSPARC-2 and display the resolution according to the gold-standard Fourier Shell Correlation criterion (GSFSC) using various automatically applied masks.

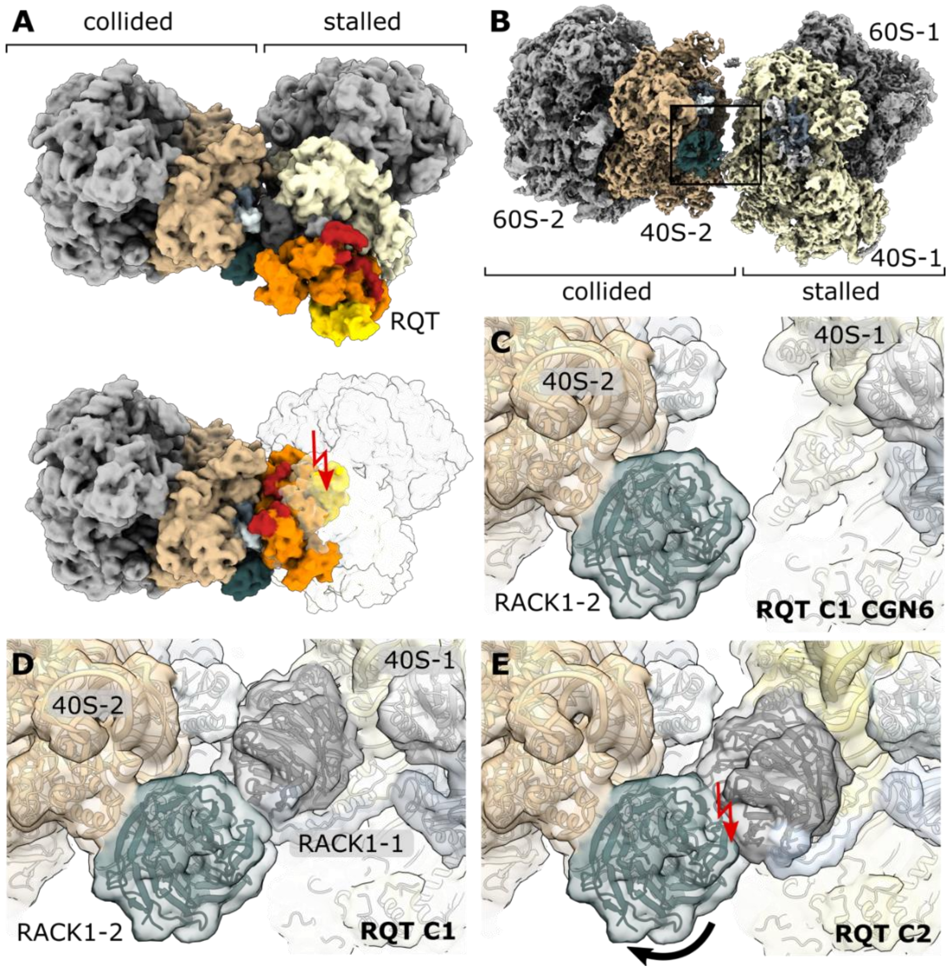


**Fig. S7: Position of RQT bound to disomes**

(**A**) RQT-bound to the lead ribosome of the disome unit (our structure) compared to a hypothetical disome with RQT bound to the colliding ribosome. Note that in such an arrangement RQT would sterically clash into the lead ribosome. (**B**) Structure (composite map) of the $(CGN)_6$-stalled disome (short overhang); view focusing on the RACK1 mediated disome interface. Box indicates zoomed region shown in (**C**). Zoom view with the model of the $(CGN)_6$-stalled disome. Note the absence of RACK1 in the lead ribosome. (**D**) view as in (**C**) displaying the canonical RACK1-RACK1 interface in a disome. (**E**) Superposition of the RQT-80S in C2 (RQT C1) conformation with a canonical disome leads to a clash between the RACK1 proteins.

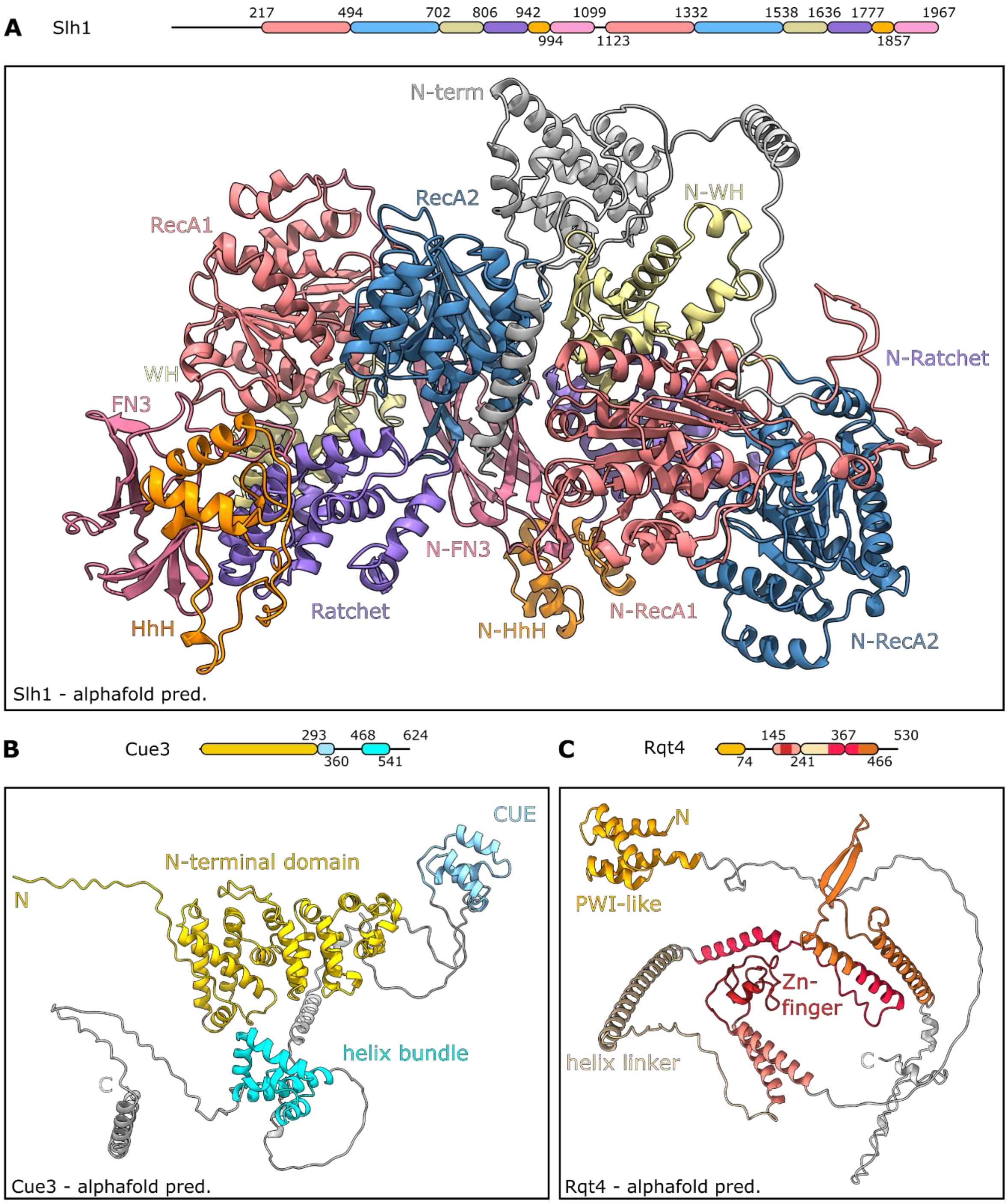


**Fig. S8: AlphaFold-2 models for RQT components**

Alphafold-2 models for Slh1 (**A**), Cue3 (**B**) and Rqt4 (**C**). Schematic representations of the domain arrangements are shown above. Density is visible in the cryo-EM map for entire Slh1 with exception of the N-terminal domain (gray), for the Cue3 N-terminal domain (yellow), the Rqt4 zinc finger (red), the C-terminal part of the Rqt4 helix linker, the following extended stretch and a part of the first helix of the following domain (magenta).

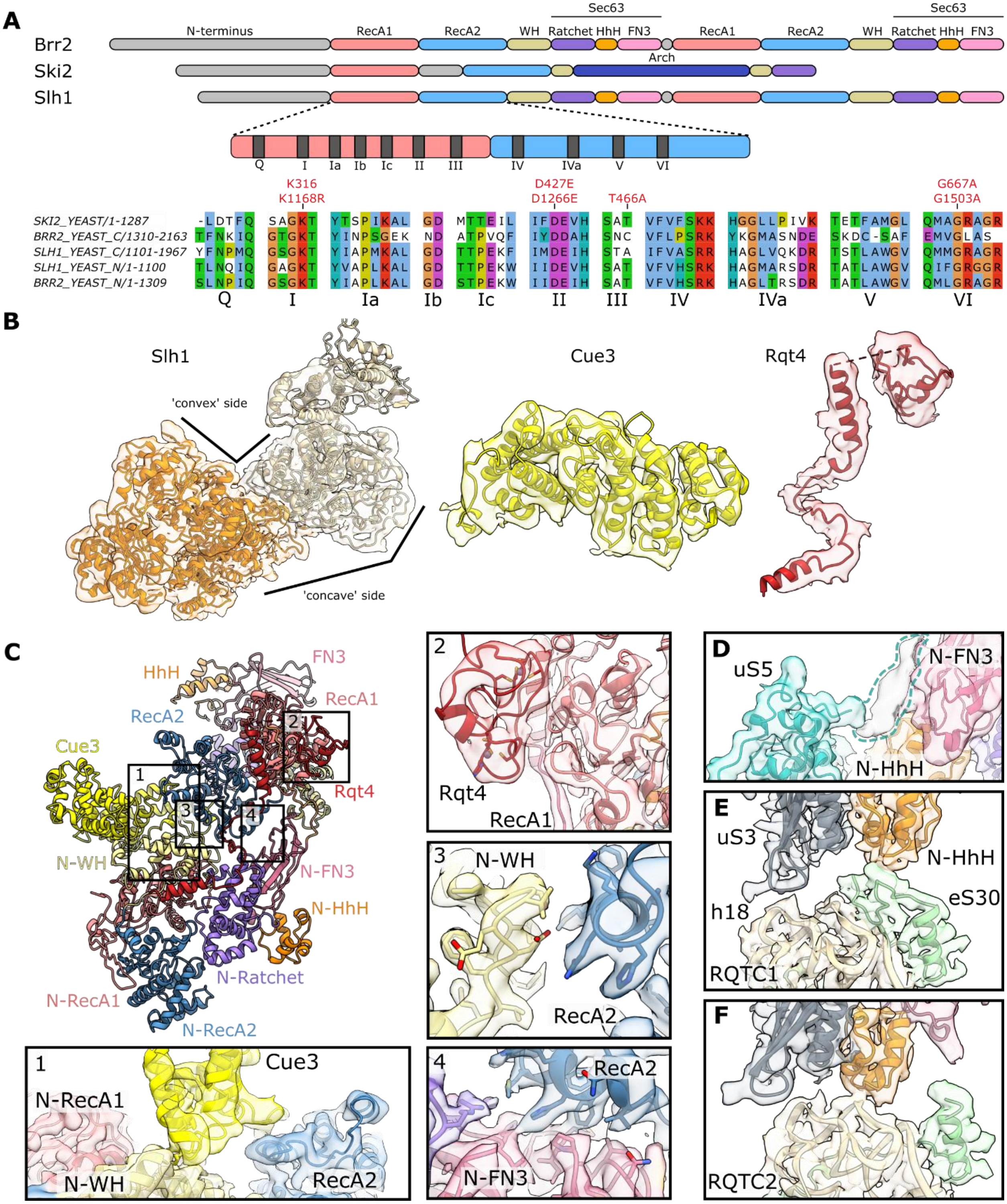


**Fig. S9: Sequence alignments and model-to-map fits for the RQT complex**

(**A**) Domain overview (top) and sequence alignments (bottom) of Slh1 with Brr2 and Ski2 from *S. cerevisiae*. The region of the (N-terminal) RecA domains is enlarged and conserved hallmark sequences for this protein family are indicated, for which also the sequence alignment is shown. Alignments are comparing the C- versus N-terminal cassettes of both Slh1 and Brr2 with to Ski2.

Note, that the CTC of Brr2 in is inactive as indicated by low conservation; for the CTC of Slh1, however, all regions crucial for ATPase activity are conserved. (**B**) Fits of the AlphaFold-2 for Slh1, Cue3 and Rqt4 models into the respective densities (transparent; low-pass filtered according to local resolution). (**C**) Close-up views showing model-to-map fits for the highlighted regions (1-4). 1: Interaction of the Cue3 N-terminal (helical) domain with Slh1; 2: Interaction of the Rqt4 ZnFD with Slh1; 3 and 4: Intercassette interactions between Slh1 N-terminal winged-helix domain (3) and FN3 domain (4) with the RecA domain of the C-terminal cassette. Most interactions are established between hydrophobic amino acids (hydrophobic patch). (**D**) Close-up view on unassigned extra density between HhH and FN3 domains of the N-terminal Slh1 cassette. This density most likely attributes to the otherwise flexible N-terminal tail of uS5. (**E, F**) Close-up view on the Slh1 N-terminal cassette interactions with rRNA helix h18 and eS30. In the C1-state (**E**), the C-terminal tail of eS30 is structured and clamped between the HhH domain and h18. In the C2 state (**F**), the eS30 tail is delocalized and the HhH domain interacts directly with h18.

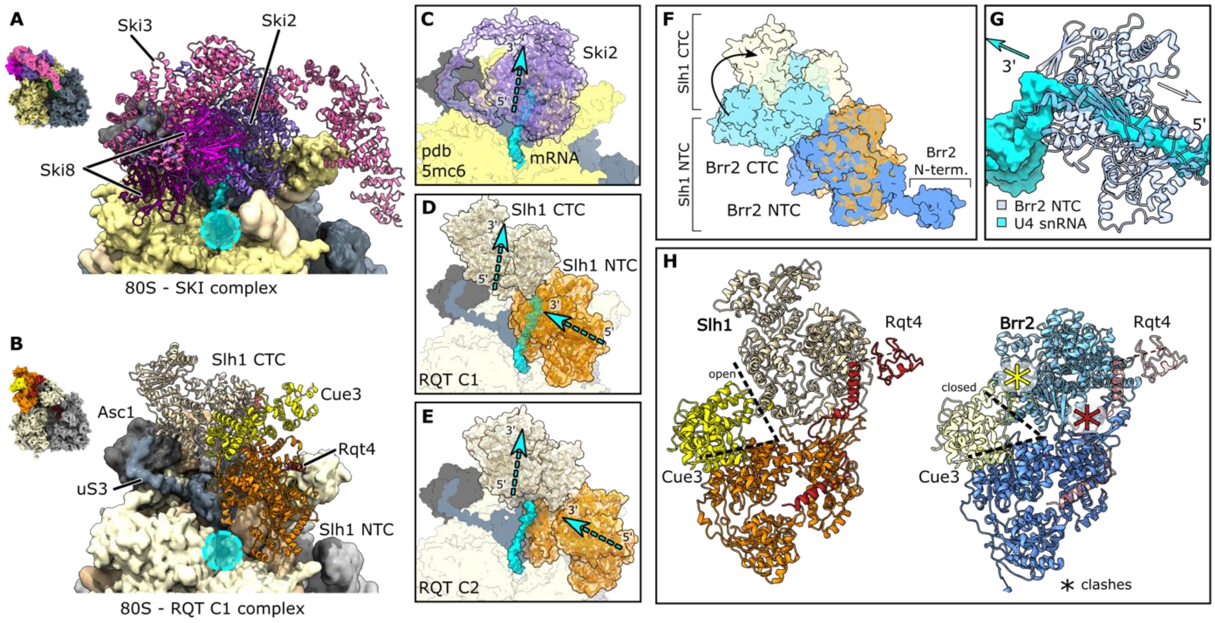


**Fig. S10: Structural comparison between RQT complex, SKI complex and Brr2**

(**A, B**) Structure of the ribosome-bound SKI complex (**A**, PDB code 5mc6) (*22*) with mRNA (cyan) emerging from the mRNA entry site (cyan circle) compared to ribosome-bound RQT complex in C1 state (**B**, this study). Thumbnails indicate the orientation. (**C-E**), Comparison of ribsome-Ski2 mRNA (**C**; cyan in all panels) after alignment of 40S subunits from RQT-80S-C1 (**D**) and RQT-80S-C2 (**D**). Note that mRNA as bound to Ski2 would clash with Slh1-NTC in C1 position, but would directly feed into Slh1-CTC in C2 position. (**F**) Superposition of ribosome-bound Slh1 with Brr2 as present in the yeast spliceosomal U4/U5.U6 tri-snRNP (*35*) (PDB code 5gan). Note that Slh1 adopts a more elongated shape. (**G**) Close-up view on the N-terminal cassette of Brr2 from the U4/U5.U6 tri-snRNP. Bound single-stranded RNA is shown in space-fill representation. Arrows indicate the direction of movement. Cyan: 5'-3' movement of the RNA, white: 3'-5'movement of the helicase. (**H**) comparison of Slh1 in the RQT complex with Brr2 from the U4/U5.U6 tri-snRNP. Note that Cue3 and Rqt4 stabilize the open Slh1 conformation and would clash with Brr2.

**Table S1. Data collection, refinement and model statistics**

| | RQTC1 (EMDB-xxxxx) (PDB-xxxx) | RQTC2 (EMDB-xxxxx) (PDB-xxxx) | 60S (EMDB-xxxxx) (PDB-xxxx) | RQTC1 stalled (EMDB-xxxxx) (PDB-xxxx) | Collided (EMDB-xxxxx) (PDB-xxxx) |
|---|---|---|---|---|---|
| **Data collection and processing** | | | | | |
| Camera | Gatan K2 summit | Gatan K2 summit | Gatan K2 summit | Gatan K2 summit | Gatan K2 summit |
| Magnification | 130.000 | 130.000 | 130.000 | 130.000 | 130.000 |
| Voltage [kV] | 300 | 300 | 300 | 300 | 300 |
| Electron exposure [e- Å $^{-2}$] | 43.6 | 43.6 | 46.4 | 44.0 | 44.0 |
| Defocus range [µm] | 0.5-3.0 | 0.5-3.0 | 0.5-3.0 | 0.5-3.0 | 0.5-3.0 |
| Pixel size [Å] | 1.045 | 1.045 | 1.045 | 1.045 | 1.045 |
| Symetry imposed | C1 | C1 | C1 | C1 | C1 |
| Micrographs collected [no.] | 21.171 | 21.171 | 14.092 | 16.508 | 16.508 |
| Initial particle images [no.] | 2.415.630 | 2.415.630 | 304.209 | 2.129.558 | 2.129.558 |
| Final particle images [no.] | 194.186 | 20.380 | 25.072 | 17.885 | 95.228 |
| Map resolution (80s) | 2.4 | 3.0 | 3.2 | 3.2 | 2.5 |
| FSC theshold | 0.143 | 0.143 | 0.143 | 0.143 | 0.143 |
| Map resolution (RQT) | 3.5 | 4.8 | | 4.3 | |
| **Refinement** | | | | | |
| initial model used [PDB code] | 6snt, alphafold | 6snt, alphafold | 6hd7, 6snt, 1G62 | 6snt, alphafold | 6snt,6i7o |
| Model resolution | 2.85 | 2.98 | 3.42 | 3.16 | 2.93 |
| FSC threshold | 0.5 | 0.5 | 0.5 | 0.5 | 0.5 |
| Map sharpening Bfactor | -66 | -40 | -10 | -38 | -59 |
| Model composition | | | | | |
| Non-hydrogen atoms | 218.512 | 218.376 | 127.849 | 218.512 | 203.400 |
| Protein residues | 13.293 | 13.289 | 6416 | 13.293 | 11.140 |
| Nucleotide residues | 5310 | 5309 | 3603 | 5310 | 5414 |
| Ligands | 95 | 13 | 24 | 95 | 93 |
| R.m.s deviations | | | | | |
| Bond lengths[Å] | 0.003 | 0.004 | 0.002 | 0.009 | 0.009 |
| Bond angles[°] | 0.569 | 0.594 | 0.534 | 0.823 | 0.732 |
| Validation | | | | | |
| Molprobity score | 1.74 | 1.75 | 1.76 | 2.05 | 1.92 |
| Clash score | 8.73 | 9.42 | 12.25 | 15.09 | 12.87 |
| Poor rotamers [%] | 0.03 | 0.03 | 0.02 | 0.05 | 0.03 |
| Ramachandran plot | | | | | |
| Favored | 96.10 | 96.21 | 97.12 | 94.74 | 95.72 |
| Allowed | 3.69 | 3.57 | 2.50 | 5.04 | 4.07 |
| Disallowed | 0.21 | 0.21 | 0.38 | 0.21 | 0.21 |

### Table S2. Strains and Plasmids

| **Yeast strains** | **Genotype** | **Reference** |
|---|---|---|
| ltn1Δ | MATa ade2 his3 leu2 trp1 ura3 can1 ltn1Δ::kanMX4 | (*61*) |
| slh1Δltn1Δ | MATa ade2 his3 leu2 trp1 ura3 can1 slh1Δ::natMX4 ltn1Δ::kanMX4 | (*7*) |
| ski2Δ uS10-3HA | MATa ade2 his3 leu2 trp1 ura3 can1 ski2Δ::kanMX4 uS10-3HA::HISMX6 | (*8*) |
| xrn1Δ, slh1Δ, cue2Δ | MATa his3Δ1 leu2Δ0 met15Δ0 ura3Δ0 can1Δ::STE2pr-his5+ lyp1Δ xrn1Δ::kanMX cue2::NatMX4 slh1::hphMX6 ade2::RFP_Pgal_GFP-2A-FLHIS3-CGA12-HIS3_MET17 | (*62*) |
| **Plasmids** | | |
| p415GPD-Hel2-Flag | CEN, LEU2, GPD promoter, HEL2-FLAG | (*7*) |
| p425GAL-Rqt2-FTP | 2μ, LEU2, GAL1 promoter, RQT2-FTP | (*8*) |
| p425GAL-Rqt2K316R-FTP | 2μ, LEU2, GAL1 promoter, RQT2K316R-FTP | (*8*) |
| p424GAL-Rqt3 | 2μ, TRP1, GAL1 promoter, RQT3 | (*8*) |
| p426GAL-Rqt4 | 2μ, URA3, GAL1 promoter, RQT4 | (*8*) |
| p7XC3GH-eIF6 | pET26b, CmR, KanR, T7 promoter,eIF6-3D-GFP-10xHis | (*45*) |
| pGEX-UBC4 | pGEX, AmpR, tac promoter, GST-3C-UBC4 | (*46*) |
| pEX-His-v5-Rpl4a-CGN12 | pEX, AmpR, T7 promoter, 6xHis-V5-TEV-Rpl4A 4-64-CGN12 | This study |
| p415-Rqt2 | CEN, LEU2, SLH1 promoter, RQT2 | This study |
| p415-Rqt2K316R | CEN, LEU2, SLH1 promoter, RQT2K316R | This study |
| p415-Rqt2D427E | CEN, LEU2, SLH1 promoter, RQT2D427E | This study |
| p415-Rqt2T466A | CEN, LEU2, SLH1 promoter, RQT2T466A | This study |
| p415-Rqt2G667A | CEN, LEU2, SLH1 promoter, RQT2G667A | This study |
| p415-Rqt2K1168R | CEN, LEU2, SLH1 promoter, RQT2K1668R | This study |
| p415-Rqt2D1266E | CEN, LEU2, SLH1 promoter, RQT2D1266E | This study |
| p415-Rqt2G1503A | CEN, LEU2, SLH1 promoter, RQT2G1503A | This study |
| p415-Rqt2K316/1168R | CEN, LEU2, SLH1 promoter, RQT2K316/1168R | This study |
| p415-Rqt2D427/1266E | CEN, LEU2, SLH1 promoter, RQT2D427/1266E | This study |
| p415-Rqt2G667/1503A | CEN, LEU2, SLH1 promoter, RQT2G667/1503A | This study |
| p425GPD-Rqt2-FTP | 2μ, LEU2, GPD promoter, RQT2-FTP | This study |
| p425GPD-Rqt2K316R-FTP | 2μ, LEU2, GPD promoter, RQT2K316R-FTP | This study |
| p425GPD-Rqt2D427E-FTP | 2μ, LEU2, GPD promoter, RQT2D427E-FTP | This study |
| p425GPD-Rqt2T466A-FTP | 2μ, LEU2, GPD promoter, RQT2T466A-FTP | This study |
| p425GPD-Rqt2G667A-FTP | 2μ, LEU2, GPD promoter, RQT2G667A-FTP | This study |
| p425GPD-Rqt2K1168R-FTP | 2μ, LEU2, GPD promoter, RQT2K1168R-FTP | This study |
| p425GPD-Rqt2D1266E-FTP | 2μ, LEU2, GPD promoter, RQT2D1266E-FTP | This study |
| p425GPD-Rqt2G1503A-FTP | 2μ, LEU2, GPD promoter, RQT2G1503A-FTP | This study |
| p425GPD-Rqt2K316/1168R-FTP | 2μ, LEU2, GPD promoter, RQT2K316/1168R-FTP | This study |
| p425GPD-Rqt2D427/1266E-FTP | 2μ, LEU2, GPD promoter, RQT2D427/1266E-FTP | This study |
| p425GPD-Rqt2G667/1503A-FTP | 2μ, LEU2, GPD promoter, RQT2G667/1503A-FTP | This study |
| p416GPD-GFP-R(CGN)12-FLAG-HIS3 | CEN, URA3, GPDp-GFP-R(CGN)12-FLAG-HIS3 | (*63*) |

**Movie S1.**

Clearing of ribosome collisions by RQT via subunit dissociation.
